# Rapid, open-source, and automated quantification of the head twitch response in C57BL/6J mice using DeepLabCut and Simple Behavioral Analysis

**DOI:** 10.1101/2025.04.28.650242

**Authors:** Alexander D. Maitland, Nicholas R. Gonzalez, Donna Walther, Francisco Pereira, Michael H. Baumann, Grant C. Glatfelter

**Author notes:** Corresponding Author* **Grant C. Glatfelter** - Designer Drug Research Unit, Intramural Research Program, National Institute on Drug Abuse, Baltimore, Maryland 21224, United States.

## Abstract

Serotonergic psychedelics induce the head twitch response (HTR) in mice, an index of serotonin (5-HT) 2A receptor (5-HT_2A_) agonism and a behavioral proxy for psychedelic effects in humans. Existing methods for detecting HTRs include time-consuming visual scoring, magnetometer-based approaches, and analysis of videos using semi-automated commercial software. Here, we present a new automated approach for quantifying HTRs from experimental videos using the open-source machine learning-based toolkits, DeepLabCut (DLC) and Simple Behavioral Analysis (SimBA). Pose estimation DLC models were trained to predict X,Y coordinates of 13 body parts of C57BL/6J mice using historical experimental videos of HTRs induced by various psychedelic drugs. Next, a non-overlapping set of historical experimental videos was analyzed and used to train SimBA random forest behavioral classifiers to predict the presence of the HTR. The DLC+SimBA approach was then validated using a separate subset of visually scored videos. DLC+SimBA model performance was assessed at different video resolutions (50%, 25%, 12.5%) and frame rates (120, 60, 30 frames per second or fps). Our results indicate that HTRs can be quantified accurately at 50% resolution and 120 fps (precision = 95.45, recall = 95.56, F_1_ = 95.51) or at lower frame rates and resolutions (i.e., 50% resolution and 60 fps). The best performing DLC+SimBA model combination was deployed to evaluate the effects of bufotenine, a tryptamine derivative with uncharacterized potency and efficacy in the HTR paradigm. Interestingly, bufotenine only induced elevated HTRs (ED_50_ = 0.99 mg/kg, max counts = 24) when serotonin 1A receptors (5-HT_1A_) were pharmacologically blocked and activity at other sites of action may also impact its pharmacological effects (e.g., serotonin transporter). HTR counts for a subset of 21 videos from bufotenine experiments were strongly correlated for DLC+SimBA vs. visual scoring and semi-automated software detection methods (*r* = 0.98 and 0.99). Finally, the DLC+SimBA approach displayed high accuracy when compared to visual scoring of HTRs for three serotonergic psychedelic drugs with variable HTR frequencies (*r* = 0.99 vs. mean visual scores from 3 blinded raters). In summary, the DLC+SimBA approach represents a modular, noninvasive, and open-source method of HTR detection from experimental videos with accuracy comparable to magnetometer-based approaches and greater speed than visual scoring.

---

Serotonin (5-HT) 2A receptor (5-HT_2A_) agonists that induce psychedelic subjective effects in humans induce the head-twitch response (HTR) in mice, a rapid paroxysmal movement of the head.*^1–4^* The HTR is a reliable behavioral proxy of psychedelic drug effects in humans, with a strong positive correlation between drug potencies for HTR induction in C57BL/6J mice and doses inducing subjective psychedelic activity in humans.*^5^* While humans do not display HTRs in response to psychedelics and some false negative and positives have been reported, measuring the HTR is a translational, accessible, replicable, and widely used preclinical paradigm used to predict psychoactivity of 5-HT_2A_ agonists.*^3, 5^* By providing a simple and reliable quantitative predictor of psychedelic-like drug effects, the mouse HTR has translational utility for examining the structure-activity relationships of serotonergic psychedelic compounds.*^3, 5–11^*

Existing approaches for quantifying HTRs are effective but have limitations. Visual scoring of experimental video recordings is the simplest and most common approach for detecting HTRs and is often considered the “gold standard” for assessing accuracy of other methods of detection.*^3, 12–17^* However, visual scoring is time-consuming and requires multiple trained raters due to inter-individual variability in scoring and potential bias introduced by visual review. The HTR can also be quantified using magnetometer-based approaches, which involve detection of changes in coil voltage caused by a magnetic reporter surgically implanted on the skull or tags clipped on the ears.*^1, 18–20^* Magnetometer-based approaches allow identification of characteristic HTR wave forms facilitating rapid quantification of HTRs within minutes of completion of an experiment.*^1, 19–21^* More recently, automated methods employed machine learning (ML) approaches to extract HTRs from magnetometer recordings and further improved detection.*^22^*

Despite the high accuracy and rapid data collection of magnetometer-based approaches to measure HTR events, this strategy has important caveats. First, current magnetometer-based methods require either surgical implantation of magnets on the skull or brief anesthesia for attachment of magnets to the ears.*^1, 5, 19, 21, 23^* Second, magnetometer setups are difficult to standardize given that the magnitude of the response depends on the physical dimensions of the coil and magnet size.*^1, 24^* A recent deployment and validation of the magnetometer-based detection system by Nakamura et al. *^25^* demonstrated high HTR detection accuracy (99%), but validation testing revealed a mean error detection rate of 11.20%. Third, the measurement of other behaviors can be impeded by using magnetometer coil chambers and the magnets themselves may interfere with running other desired in vivo assays. Lastly, novel compounds may produce unforeseen behaviors that increase false positive detection using magnetometer approaches.*^19, 21, 22^* The unpredictable behavioral effects of new pharmacological agents can be mitigated more easily with video-based approaches which allow for manual review or assessment of behavioral effects of each drug.

Video-based HTR detection in mice has been reported using several approaches.*^1, 26^* Our group has previously validated a semi-automated, commercial software-based method to detect HTRs from video recordings. *^26^* This approach allows for rapid manual review of each time-stamped HTR event, but high-throughput drug screening is challenging due to lack of automaticity and scalability. Furthermore, commercial software licenses may be prohibitively expensive for some laboratories. Therefore, there is a need to develop rapid, non-invasive, open-source, accurate, and automated HTR detection approaches based on the analysis of experimental video recordings. ML-based analyses of rodent behavior outperform commercial software systems with human-level accuracy*^27^* and may present an improvement upon existing video-based HTR tracking strategies. In parallel to visual scoring, ML approaches have been reported for HTR detection in mice and to quantify HTR-like wet-dog shakes in rats.*^28, 29^* However, the existing methods are limited in scope, detail, and validation assessments necessary for deployment in a drug discovery context.

To address these challenges, we developed a novel approach to characterize the HTR in mice using the open-source ML toolkits, DeepLabCut (DLC) and Simple Behavioral Analysis (SimBA). Using random historical experimental videos featuring HTR events across psychedelic chemotypes and experimental conditions, pose estimation data were extracted using DLC.*^30, 31^* The analysis time required to extract pose estimation data is a function of both the video resolution (i.e., the level of detail in pixels) and its frame rate (i.e., the frequency of images captured per unit of time). To allow evaluation of this trade-off, multiple models were trained using different video resolutions and frame rates to compare body part tracking performance and analysis speed. The experimental videos were downscaled from the original resolution with a frame height and width of 1280×960 pixels to 50% (906×680), 25% (640×480), and 12.5% (454×340) video resolutions, and frames rates from 120 frames per second (fps) to 60 and 30 fps. SimBA was then used to generate behavioral classifiers for the HTR for each combination of resolution and frame rate. Each DLC+SimBA model combination was then applied to a subset of 38 historical experimental videos, not used in training of DLC or SimBA models, to evaluate performance. To demonstrate the utility of the approach in psychedelic drug discovery, the best performing DLC+SimBA model combination was deployed to characterize bufotenine (5-hydroxy-*N*,*N*-dimethyltryptamine, 5-HO-DMT, or dimethylserotonin), a tryptamine derivative with purported psychedelic effects *^32–36^* and unknown potency and efficacy in the HTR paradigm. The HTR counts from three standard psychedelics with variable HTR frequencies were also examined in follow up deployment studies to further validate HTR detection vs. visual scoring by three independent raters. Overall, the present data show that our ML-based HTR detection tool is rapid, accurate, and can be used to create high-throughput workflows for preclinical assessment of psychedelic-like effects of compounds.

## Results and Discussion

### 2.1 Mouse pose estimation model

Our previous studies with psychedelic drug administration in mice provided a rich library of historical videos depicting HTR events induced by various drug chemotypes under differing experimental conditions.*^6, 7, 11, 26, 37, 38^* This video library served as a source for model training, testing, and validation. For body part tracking of C57BL/6J mice, we used DLC *^30, 31^* and trained a model at each video resolution (50%, 25%, 12.5% of original video resolution). The 13 tracked body parts used for each pose estimation model are shown in **Figure 1A**.

**Figure. 1.**
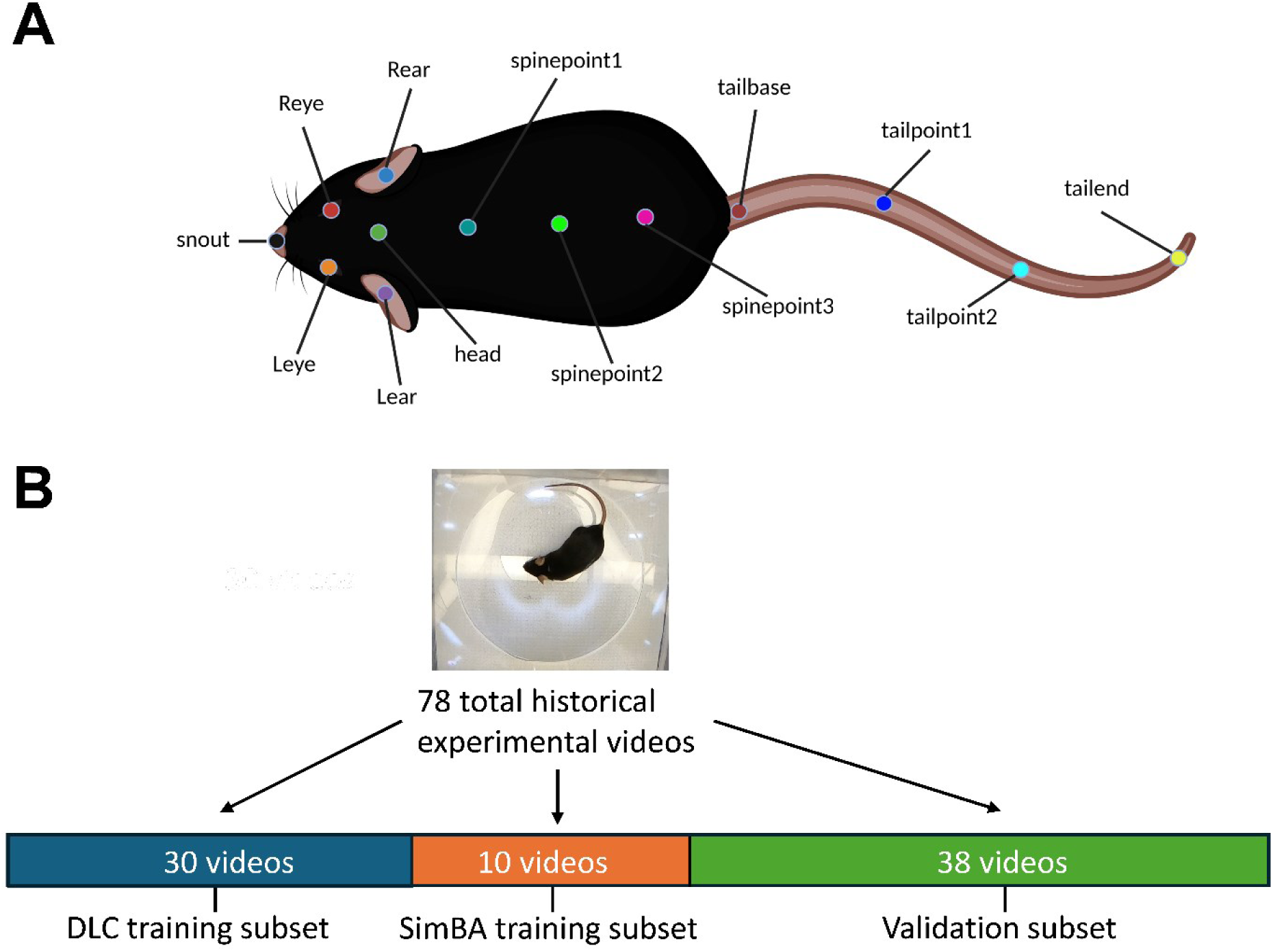
C57BL/6J mouse body parts labeled for tracking (**A**) and schematic for processing of historical videos for HTR (**B**). All pose estimation models were trained to track the 13 body parts shown. Image for panel A was created in https://BioRender.com.

Specifically, extracted frames taken from 30 videos (at each resolution) and 16 animals (**Figure 1B**) were labeled. The final snapshot for each DLC model (computed root mean square error between the model-predicted labels and user labels) are shown in **Table S1**. Since each model was trained on the same set of videos, but at different resolutions, these are also reported as a percentage of the animal’s body length to allow a direct comparison.

These percentages suggest that the final DLC models trained at each video resolution were uniform in their body part tracking performance. Time costs for DLC model video analysis and FFmpeg video resolution downscaling for a subset of 10 experimental videos are reported in **Table S2**. As expected, the time costs for using FFmpeg and DLC decreased in a frame rate- and resolution-dependent manner (**Table S2**). Time costs for DLC pose estimation model training are reported in **Table S3.** Importantly, as recommended by Mathis et al., the training process was repeated to optimize each model’s performance, requiring analysis of additional videos to extract video frames with poor model labeling and re-training the model. The total time cost to repeat the training process 3 times for the final 50% pose estimation model in DLC was about 2,848.91 minutes, or 47.40 hours. However, the majority of model training steps proceeded automatically, and the labeling or correcting of frames to improve model performance represented the only labor-intensive manual task during the training process. In total, the estimated time cost for the manually labeling or correcting 600 frames for the 50% DLC model was about 206.64 minutes, or 3.44 hours, with similar time costs for DLC models at 25% and 12.5% resolutions. Internal controls featuring an empty cylinder arena showed no animal tracking for any of the three DLC models (data not shown), and a historical experimental video featuring an immobile mouse featured consistent tracking for each of the three DLC models (data not shown). Visual inspection of labeled validation subset videos further confirmed model performance. Overall, these values highlight that although DLC pose estimation model training requires about 47 hours, only a few hours of manual labeling are needed to achieve accurate body part tracking.

### 2.2 HTR behavioral classifier evaluation

After successfully training and validating pose estimation models, SimBA was used to generate 9 behavioral classifiers from a non-overlapping subset of 10 random historical experimental videos containing 136 HTR events (**Figure 1B**). Time costs for SimBA behavioral classifier training for the 50%, 120 fps model combination are reported in **Table S4.** The total time cost for training the 50%, 120 fps behavioral classifier in SimBA was 105.91 minutes, or 1.77 hours. While most steps of behavioral classifier model training proceed automatically, labeling HTR events was manual and required 65.95 minutes to complete. Each of the 9 behavioral classifiers was then evaluated in a separate ’hold-out’ validation subset containing 38 historical experimental video recordings, to assess the generalizability of each model combination (not included in the training or test set). Each classifier was evaluated in every frame of each of 38 historical experimental video recordings in the validation video subset, generating a prediction of whether a HTR event was present (positive) or not (negative). These predictions were compared against ground truth visually scored by three independent raters (*r* = 0.92 – 0.99, *p* < 0.0001). The scoring metrics calculated were the precision (true positives/all true and false positive events), recall (true positives/all true positive and false negative events), and F_1_ scores (harmonic mean of precision and recall) across all the frames in the validation video subset (**Figure 2 & 3**).

**Figure 2.**
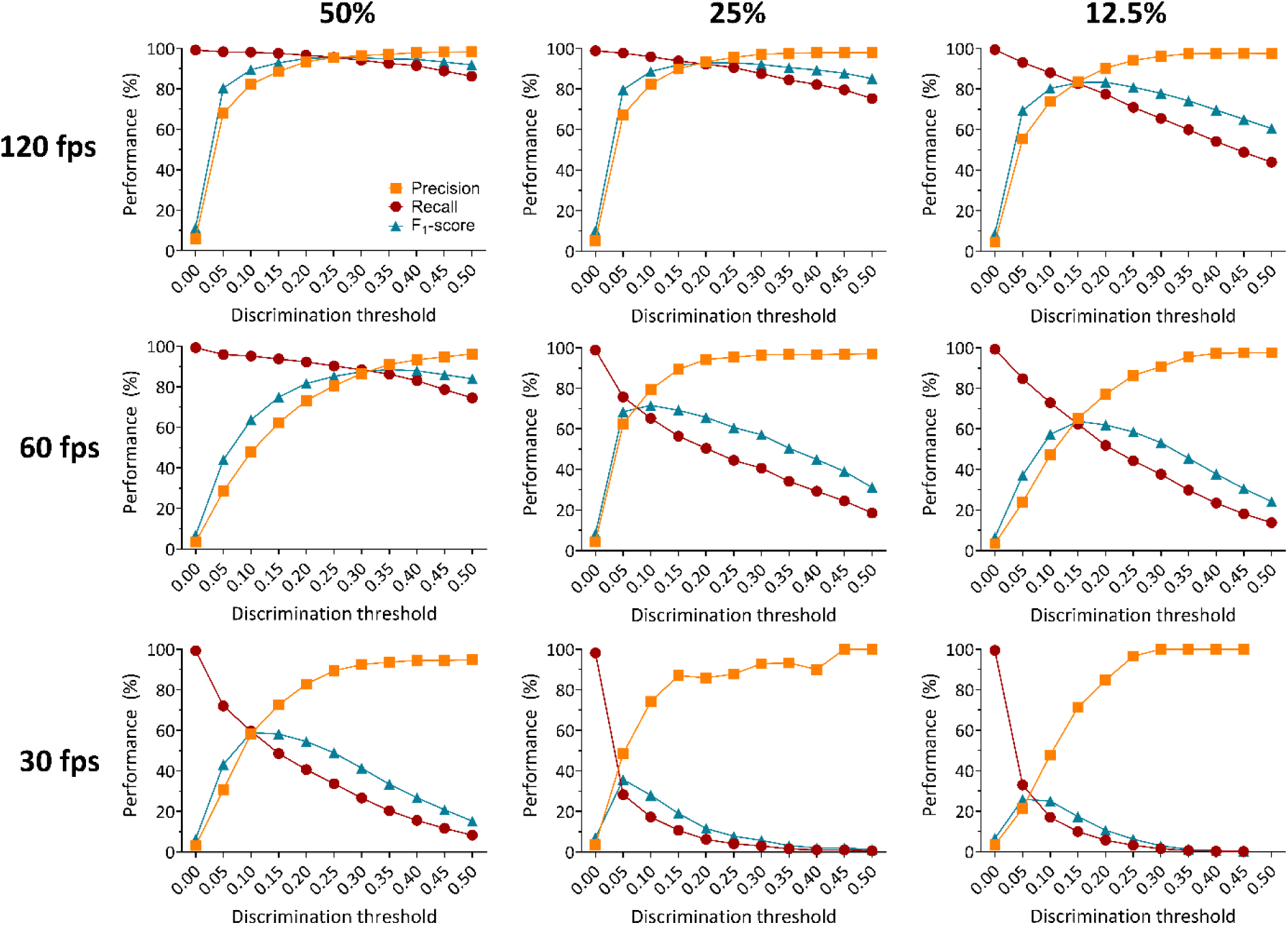
Accuracy-discrimination threshold curves for SimBA behavioral classifiers for 50%, 25%, and 12.5% video resolution at 120, 60, and 30 fps. Discrimination thresholds are the probability levels at which the behavioral classifier determines an HTR is detected. All minimum behavior bout lengths were set to 30 ms. Precision in red, recall in orange, and F_1_ scores in teal at different discrimination thresholds are shown for all behavioral classifiers.

These validation videos included 18 experimental videos from Glatfelter et al. *^26^*, and 20 additional random historical experimental videos of various psychedelic drug treatments (i.e. psilocybin, lysergic acid diethylamide or LSD, 2,5-dimethoxy-4-iodoamphetamine or DOI, 2,5-dimethoxy-4-butylamphetamine or DOBU, 2,5-dimethoxyamphetamine or 2,5-DMA, and 2,5-dimethoxy-4-propylamphetamine or DOPR), doses (0.001 – 30 mg/kg), lighting conditions (standard room overhead lights with additional accessory lights placed above the open field areas), and animals (male and female), for a total of 38 experimental videos (**Figure 1B**). Each of the 9 behavioral classifiers were assessed at various discrimination thresholds, from 0% to 50% (**Figure 2 & 3**).

**Figure 3.**
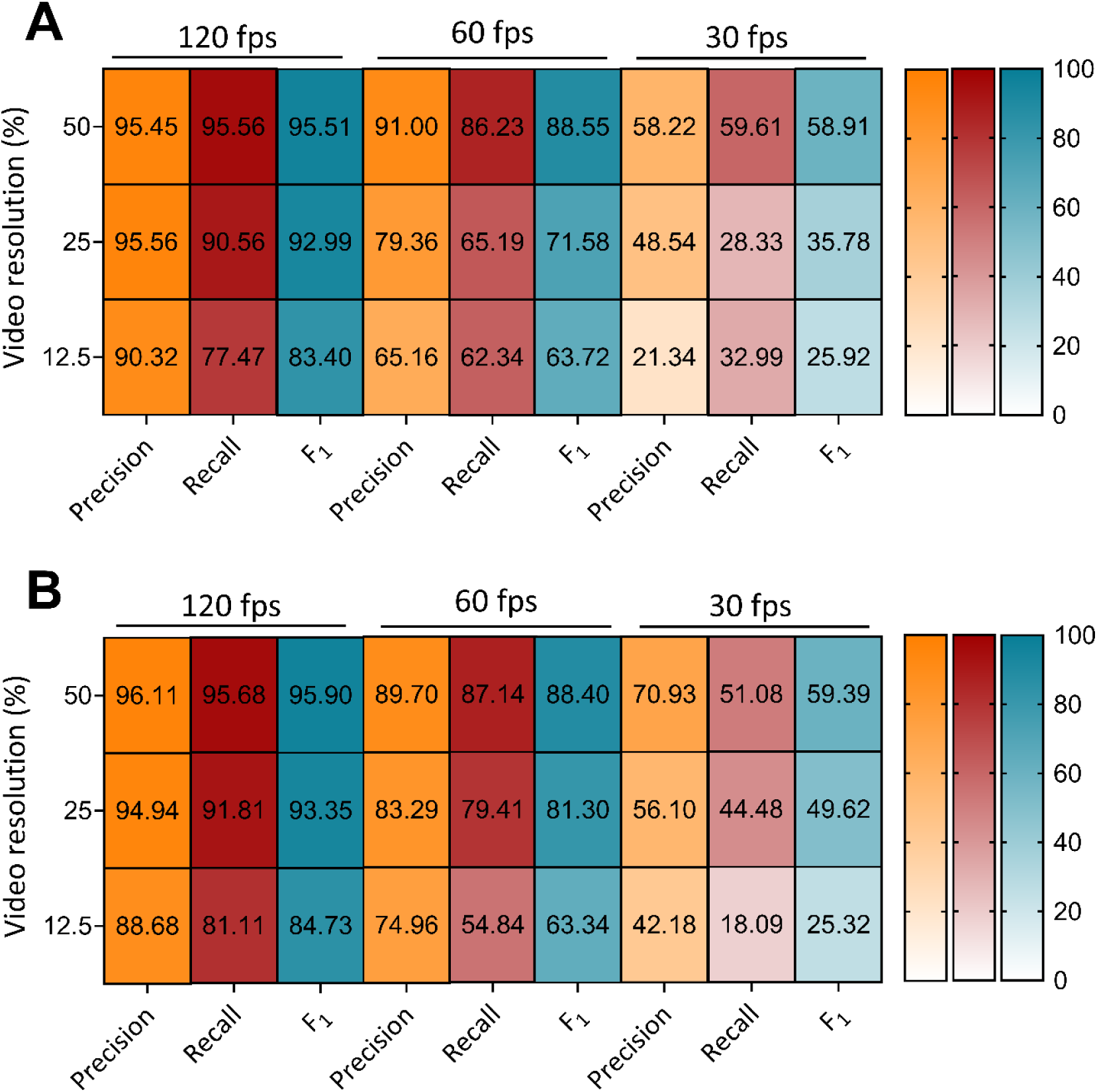
Heatmap visualization summarizing performance metrics at best performing accuracy-discrimination threshold for each SimBA classifier for 50%, 25%, and 12.5% video resolution at 120, 60, and 30 fps annotated originally (**A**) and by another experimenter (**B**). All minimum behavior bout lengths were set to 30 ms.

Visual scores of the HTRs from three independent raters indicated the validation video subset contained 879 total HTR events. HTR classifier performance on the validation video subset decreased in a frame rate and resolution dependent manner, with high performance for each respective DLC+SimBA model combination at 120 fps irrespective of video resolution (F_1_ = 95.51 - 83.40), with lower performance for 60 fps (F_1_ = 88.55 - 63.72) and 30 fps (F_1_ = 58.91 - 25.92) (**Figure 2 & 3, Figure S1**). Notably, the best performing 50% video resolution video and 60 fps model combination (F_1_ = 88.55 at 0.35 discrimination threshold) outperformed the best performing 12.5% video resolution at 120 fps model (F_1_ = 83.40 at 0.20 discrimination threshold), suggesting that high performance can be achieved even with lower resolution and frame rate videos. While not tested in the current study, training set construction in SimBA can influences F_1_ score *^39^*, and performance may further improve with more training videos containing HTR events. To account for variance in subjective annotations of HTR and further confirm the validity of these findings, we compared this performance to 9 behavioral classifiers annotated by another experimenter, which produced similar accuracy-discrimination threshold curves and values (**Figure S2, Figure S3**). The patterns of performance as measured by best F_1_ scores across DLC+SimBA model combinations were relatively indistinguishable, though optimal discrimination thresholds for HTR detection varied slightly (e.g. 0.25 to 0.30 at 50% resolution, 120 fps). Despite this, the overall performance similarities suggested that the performance of DLC+SimBA model combinations are similar between annotators.

As expected, performance was reduced as video resolution and frame rate decreased. The best performing model combination, as measured by F_1_ score, was the 50% resolution, 120 fps model at a discrimination threshold of 0.25. These results suggest that high frame rate video recordings are optimal for detecting the HTR behavior. For example, the 50%, 120 fps DLC+SimBA models at the 0.25 discrimination threshold featured a high precision (95.45) and recall (95.56), meaning that each HTR prediction had a 95% likelihood of correctly identifying a true HTR, and these models captured 96% of all true HTR events in the validation video subset. Together, these findings support the deployment of this approach for automatic quantification of HTR events in experimental videos with moderate to high frame rates (60 – 120 fps) and low to moderate scaled resolutions (25 – 50%).

### 2.3 Post-hoc explainability metrics for HTR behavioral classifiers

A behavioral classifier produced with the SimBA toolkit makes its predictions based on the values of a set of features tracked by DLC. In addition to the prediction, SimBA also provides a quantitative measure of the importance of each feature to the prediction of the HTR behavior by the classifier. For the specific classifier type used in SimBA, an ensemble model of decision trees, this is the mean decrease in impurity (also referred to as gini importance), a measure of the total decrease in node impurity averaged over all trees of the network/ensemble *^40^*. Averages for relative feature importances of each behavioral classifier at each resolution and frame rate were calculated for heatmap visualization (**Figure 4**). Based on mean decrease in impurity values, feature contributions were variable across classifiers, frame rates, and resolutions (range = 0.000428 – 0.132). This suggests that reliable tracking of some features may be affected by either of these factors, forcing classifiers to rely on other features with redundant or complementary information. Based on this metric, movement of the left ear (range = 0.0421 – 0.115), right ear (range = 0.0421 – 0.132), head (range = 0.0405 – 0.125), and first spine point labels (range = 0.0399 – 0.0994) were most important for predicting the HTR across all 120 fps and the 50%, 60 fps model combinations, with lower probability for detections among other features (range = 0.00043 – 0.0747).

**Figure 4.**
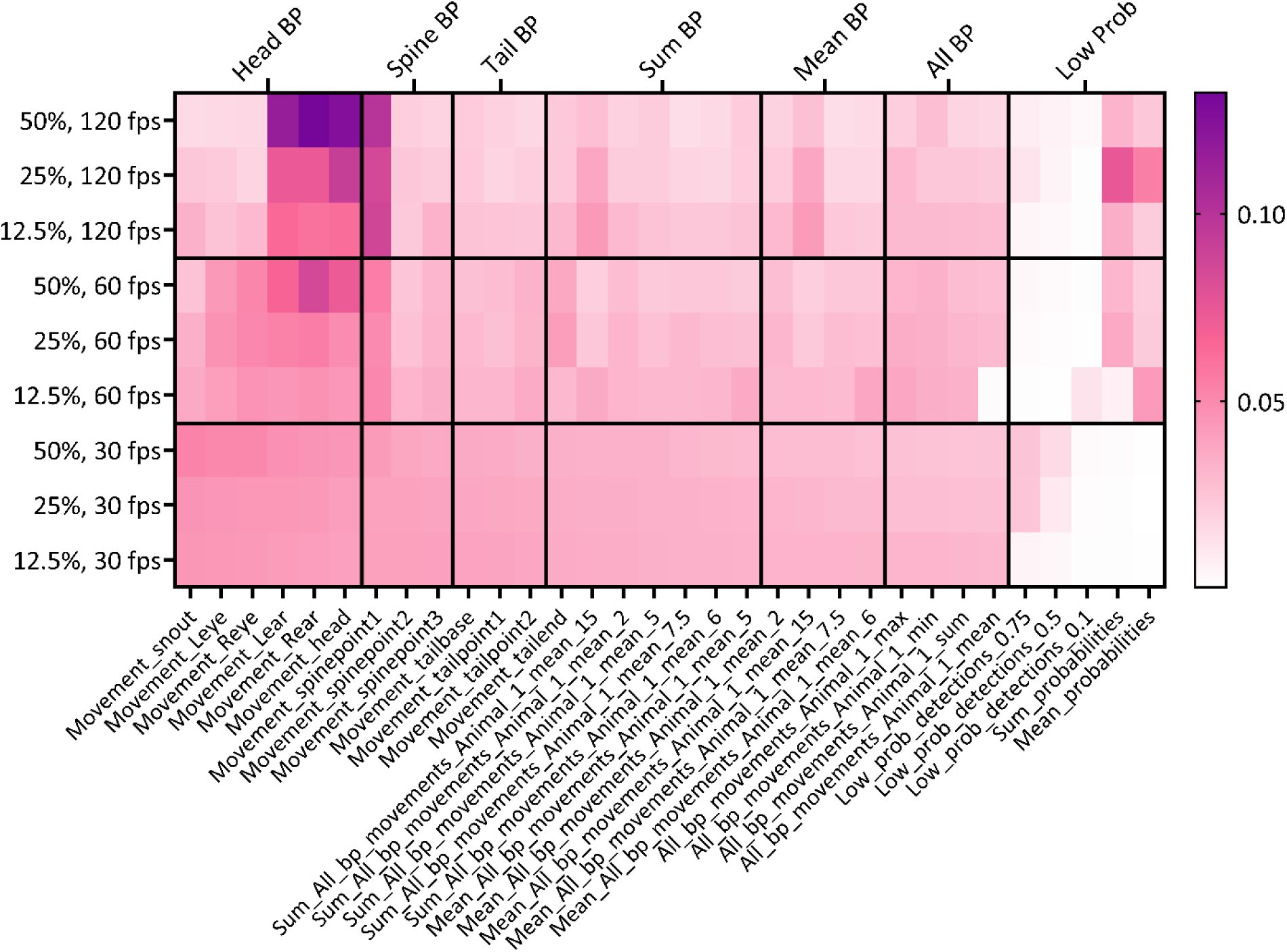
Feature importance as determined by mean decrease in impurity (gini importance) of all calculated features. The tile color corresponds to the relative importance value of each feature.

Relative importance of various categories was also visualized to further enhance interpretability by averaging related features (i.e. all features extracted from head body parts) (**Figure 4**). Head and upper-back body part movement features were the most important feature categories for the best performing model combinations across video resolutions at 120 fps as well as for the 50%, 60 fps model. Those feature categories were more important compared to the tail body part movement, all body part movement, and probabilities feature categories. The greater values for the head and upper-back features in these model combinations suggests a strong impact on predicting the HTR.

The feature importance indicates how much each feature contributes to the classifier prediction on average, across all the data that it was trained on. Beyond this, users may be interested in an explanation of which features contributed the most for the probability assigned to each behavior in a particular frame. For this purpose, SimBA provides Shapley Additive Explanations (SHAP), an explainability metric based on a game theoretic approach,*^41^* which is particularly suitable for the random forest classifiers SimBA produces.

SHAP analyses were conducted in SimBA on 100 random frames with HTR behavior present and 100 random frames with HTR behavior absent for each classifier, to measure the individual feature contributions to the overall frame behavior probability. In agreement with results using mean decrease in impurity, the Pearson correlations between each SHAP value sampled from frames where the behavior is present and absent show that features related to head and upper-back seem to substantially impact on HTR behavior probability for all resolutions at 120 fps as well as the 50%, 60 fps model (data not shown). For both post-hoc explainability metrics, the high importance of head and upper back-related body part movement features suggests these behavioral classifiers detect the HTR using body parts directly relevant to the response.

### 2.4 Behavioral control experiments using DLC+SimBA approach with SKF 38393-induced grooming and amphetamine-induced hyperlocomotion

The potential for false positive counts induced by competing behaviors (e.g. grooming, motor activity, jumping) are traditionally assessed in HTR detection method validation studies.*^1, 19, 21, 22, 26^* To further validate the DLC+SimBA approach, and compare present results to traditional HTR detection studies, the best performing model combination was tested for its ability to distinguish the HTR from grooming and hyperlocomotor activity.

The best performing models for HTR detection (50% resolution, 120 fps, F_1_ = 95.51) were also deployed on downscaled historical experimental videos from Glatfelter et al. *^26^* examining SKF38393-induced grooming (12 videos) and d-amphetamine induced hyperlocomotion (16 videos) for a total of 28 videos. Using historical visual HTR scores from a trained independent rater, the performance, recall, and F_1_ scores were evaluated (**Figure 5).** The behavioral classifier was assessed at various discrimination thresholds, from 0% to 95%. From these results, it was observed that the discrimination-accuracy curves were right-shifted compared to those obtained on the validation video subset. Practically, this means that higher discrimination thresholds would be required to obtain comparable measures of performance. Historical visual HTR scores indicated that these behavioral control videos contained 124 HTR events. At the best performing discrimination threshold of 0.25, the 50%, 120 fps models yielded greater false positives compared to the validation video subset evaluation (precision = 50.41) with very high recall (recall = 100). However, increasing the discrimination threshold to 60% reduced the number of false positives to levels (precision = 93.65, recall = 95.16, F_1_ = 94.4) similar to the results from in the validation subset evaluation at 25% (precision = 95.45, recall = 95.56, F_1_ = 95.51). Moreover, false positive detections decreased as the discrimination threshold increased beyond 60%, but at a greater cost to recall. Although there were few false positive detections at the 60% discrimination threshold (*n* = 8), several of these false positives occurred when the mouse turned its head while darting, or when rapidly rearing and turning its head.

**Figure 5.**
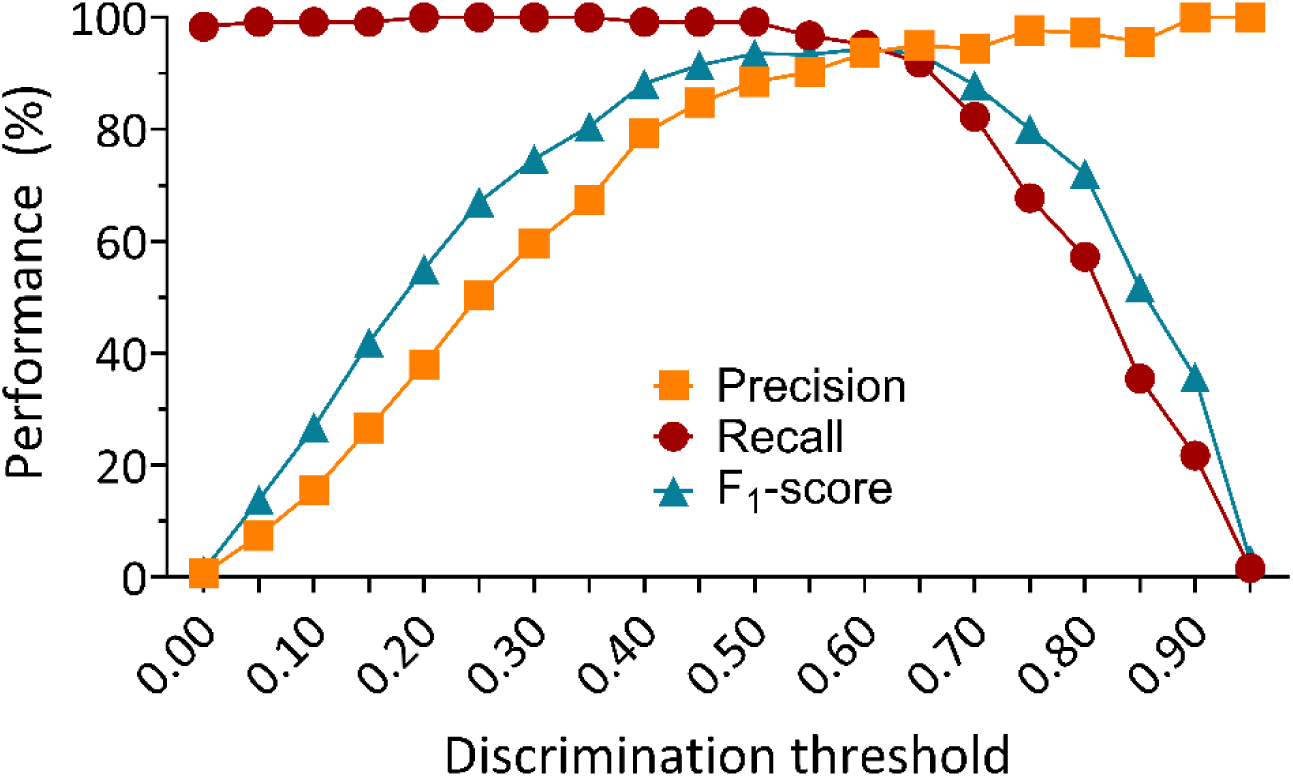
Accuracy-discrimination threshold curves using the 50% video resolution, 120 fps model combination for behavioral control videos. Discrimination thresholds are the probability levels at which the behavioral classifier determines an HTR is detected. All minimum behavior bout lengths were set to 30 ms.

Together, these findings indicate that the DLC+SimBA approach can reliably distinguish the HTR from similar behaviors, with performance equivalent to established automated HTR detection methods. By increasing the discrimination threshold, the detection of false positives was very low without sacrificing recall or precision, offering a viable strategy for addressing potential false positive counts. These findings further support the deployment of the DLC+SimBA approach for evaluating the HTR in a psychedelic drug discovery context.

### 2.5 Psychedelic-like effects of bufotenine using the DLC+SimBA HTR detection approach

To demonstrate the utility of our DLC+SimBA HTR detection approach in studying the psychedelic-like effects of 5-HT_2A_ agonists, the best performing model (50% resolution, 120 fps, F_1_ = 95.51 for the 0.25 threshold) was deployed to characterize the HTR induced by bufotenine (5-HO-DMT or *N*,*N*-dimethylserotonin, **Figure 6A**). Bufotenine is a serotonergic psychedelic nonselective for 5-HT_2A_ with uncharacterized potency and efficacy in the HTR paradigm. A recent study in mice evaluated a 1 mg/kg dose of bufotenine administered intraperitoneally, finding ∼20 – 30 HTRs *^42^*, but a full dose-response evaluation was not conducted. Here, bufotenine was administered subcutaneously (s.c.) to mice at doses ranging from 0.03 to 30 mg/kg to measure acute effects of the drug on HTR, temperature change, and locomotor activity over a 30 min testing session.

**Figure 6.**
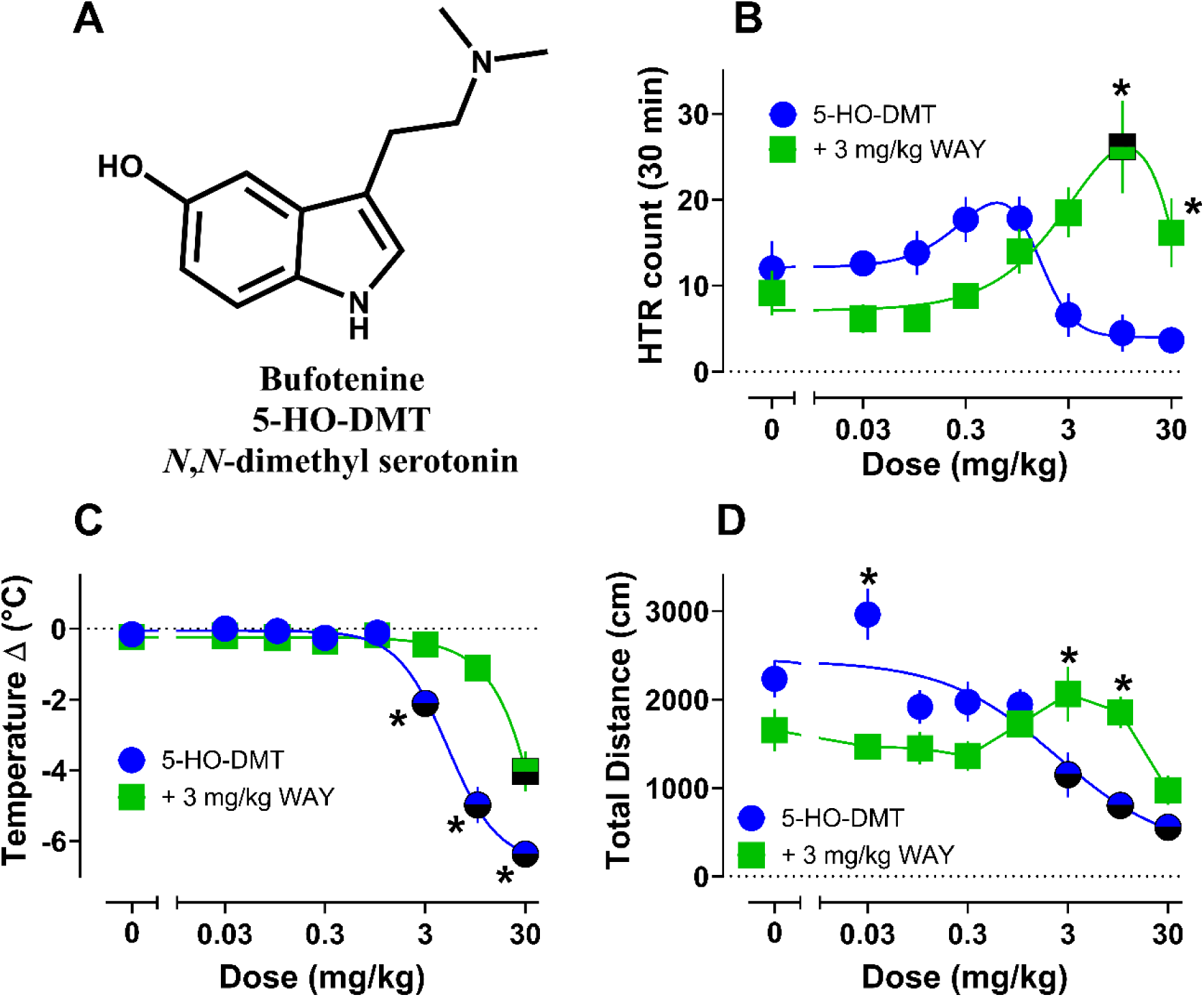
Dose-response curves for effects of bufotenine without and with WAY100635 pretreatment on HTR. (**A**) the freebase chemical structure of bufotenine. All experimental videos were analyzed using the 50%, 120 fps model DLC+SimBA combination at 0.25 discrimination threshold for HTR counts (**B**), body temperature change pre vs post session (**C**), and total distance traveled during the 30 min testing period (**D**). All values are mean ± SEM and represent *n* = 5 – 7 mice per data point. Half bolded symbols represent statistical differences from vehicle controls for each group, while asterisks represent statistical differences between groups across doses (*p* < 0.05).

Dose–response data for HTRs induced by bufotenine over the testing session are shown in **Figure 6B** while time-course data are shown in **Figure S4A**. Descriptive statistics and additional details of statistical comparisons can be found in **Table S3**. Total HTR counts over the session for bufotenine were not significantly increased versus vehicle controls across the tested doses. However, compared to vehicle controls, bufotenine did induce small increases in HTRs from 0.1 – 1 mg/kg and decreases in HTR counts from 3 – 30 mg/kg (**Figure 6B**, **Figure S4A**). The typical inverted U dose-related effect seen for related tryptamines was evident for bufotenine, but the compound was judged to be inactive given small, negligible effects on HTR counts. Importantly, the descending limb of the HTR dose-response curve corresponded with doses inducing significant hypothermic and hypolocomotor effects relative to vehicle controls (**Figure 6C & D**, **Figure S5A**, **Table S3**), in agreement with existing literature for related tryptamines.*^6, 7, 11, 15, 37, 43^*

The weak HTR activity of bufotenine observed here is notable, given that its psychedelic-like effects appear to vary depending on the study, route of administration, and dose.*^32–36, 44–47^* Bufotenine is a natural product in plants and animals that has been used as part of snuffs or extracts in various cultures for its psychoactive and other effects.*^33, 47–52^* Like other serotonergic psychedelics, bufotenine interacts with 5-HT_2A_, other 5-HT receptors (e.g. 5-HT_1A_), and non-receptor sites of action (e.g. serotonin transporter or SERT).*^53–60^* 5-HT_2A_ mediates the primary psychedelic-like effects of compounds related to bufotenine *^2, 61–63^*, but non-5-HT_2A_ activities of the drugs, such as 5-HT_1A_ and SERT, can inhibit expression of the HTR in mice and subjective effects in humans.*^6, 37, 38, 43, 64–71^*

Given that compounds with pharmacological profiles like bufotenine can be “false negatives” in the HTR paradigm *^6, 64^*, we sought to evaluate the role of 5-HT_1A_ in modulation of the HTR by the drug, especially since it displayed a biphasic dose-response curve for induction of HTR (**Figure 6B**) and induces psychedelic subjective effects in humans in some contexts.*^33, 34, 46, 48^* Mice were pretreated with the 5-HT_1A_ antagonist WAY100635 (3 mg/kg sc.) 10 mins prior to administration of bufotenine (0 – 30 mg/kg sc.), and acute effects on HTR, body temperature, and motor activity were measured for 30 min. Pretreatment with WAY100635 revealed a more pronounced HTR curve, with significantly elevated counts vs. vehicle controls at 10 mg/kg (**Figure 6B**, **Table S3**). The potency and efficacy for bufotenine to induce the HTR under these conditions (ED_50_ = 0.99 mg/kg, E_max_ = 24 counts) were similar to the results obtained with 5-methoxy-*N*,*N*-dimethyltryptamine (5-MeO-DMT) analogs and WAY100635 pretreatment.*^6^* Corresponding with the more pronounced HTR, bufotenine+WAY100635 treatment also blocked hypothermic and hypolocomotor effects except at the highest dose (30 mg/kg, **Figure 6C & D**, **Table S3**). Time-course plots for HTRs and motor activity across the 30 min session in WAY100635 pretreatment experiments are depicted in **Figure S4B** & **Figure S5B**, showing that only the highest dose had a delayed (∼10-15 min post bufotenine) effect to reduce motor activity and HTR counts.

Despite the greater HTR activity produced by bufotenine plus 5-HT_1A_ blockade, the compound was still weaker in potency and efficacy compared to related compounds such as psilocin (ED_50_ = 0.11 – 0.17 mg/kg, E_max_ = ∼23) and *N*,*N*-dimethyltryptamine (DMT; ED_50_ = 1.54 mg/kg, E_max_ = ∼76).*^5, 7^* The reduced activity of bufotenine could be due to several factors such as poor blood brain barrier penetration, rapid metabolism, route of administration, other pharmacological activities (i.e. SERT), or a combination thereof.*^33–36, 44, 47, 53, 72, 73^* Prior studies show that bufotenine is a potent substrate-type releaser at SERT *^53^*, which may impact its pharmacological effects. Here, we also studied the monoamine transporter activity of bufotenine using rat brain synaptosome preparations. Consistent with previous data *^53^*, bufotenine was a potent SERT releasing agent (SERT release EC_50_ = 97 nM) comparable to the entactogen, 3,4-methylenedioxymethamphetamine (MDMA; SERT release EC_50_ = 81 nM; **Figure S6A - C**), but the drug did not induce release at dopamine or norepinephrine transporters. The role of species differences in SERT activity (rat vs. mouse vs. human) is unclear, and may be relevant to interpretations of the present data. However, prior results examining monoamine releasing activity of tryptamines in cells transfected with human SERT vs. rat brain synaptosomes are largely in agreement.*^53, 59^* Also, given that the pharmacological effects of other serotonergic psychedelics, such as psilocybin and LSD, are influenced by SERT activity in mice and humans *^67–69, 71^*, it seems possible that SERT substrate actions of bufotenine may influence its ability to elicit the HTR.

To demonstrate the speed of our approach compared to visual HTR scoring, we timed each step of our approach using the 50%, 120 fps model combination for a representative set of 47 videos from a bufotenine experiment (**Table S5**). The time to complete this process, including downscaling to 50% video resolution, DLC inference time using the 50% resolution pose estimation model, and SimBA behavioral classifier analysis using the 50%, 120 fps model was 1,814.89 minutes, or about 30 hours. Notably, this process can be automated using simple custom Python and batch scripts. The greatest time cost was from using the 50% resolution DLC pose estimation model to analyze experimental videos (1405.70 minutes, or 23.43 hours). As evidenced by the high F_1_ scores for the 25%, 120 fps model combination,, a model combination trained using more HTR events would likely offer similar performance, halve the DLC inference time cost, and significantly reduce total time costs for HTR quantification. Training such a model would not incur significant additional time costs, given the limited time required for labeling HTR events in SimBA. Moreover, given that the time cost for visual HTR scoring is generally equal to the length of the experimental videos and requires multiple trained raters, our automated approach achieves more rapid HTR quantification than traditional visual HTR scoring. Overall, our approach represents a significant improvement in the time and resource costs necessary to quantify HTR compared to traditional visual scoring.

### 2.6 Comparing DLC+SimBA vs. visual and commercial software-based HTR detection approaches

A subset of 21 videos randomly chosen across doses/treatments from the bufotenine dose-response and WAY100635 antagonist pretreatment studies were next visually scored by an experimenter blind to drug treatments. These videos were also scored using the semi-automated commercially available approach utilizing the TopScan software platform (Clever Sys Inc.) previously validated in our laboratory.*^26^* Scores from these approaches were then compared to scores from the best performing DLC+SimBA model combination. Visual scoring results revealed that there were 326 total HTR events recorded across the 21 videos (11 bufotenine and 10 bufotenine + WAY), while the DLC+SimBA model (50%, 120 fps, 0.25 discrimination threshold) found slightly higher overall counts (331, 102% of visual) and the commercial software method found lower HTR counts (284, 87% of visual). Values from all 3 data sets were strongly correlated (**Figure 7A – C**), supporting similarity of the values across scoring methods. Importantly, the discrepancies in counts on a video-to-video level were ≤ 6 counts (mean difference between methods = ∼2), which is likely within the margin of inter-rater variability error seen with averaging visual scores from multiple raters.*^1, 74^* When manually reviewing extra events scored by the DLC+SimBA approach vs. visual scoring, it was noted that some of the scored events were missed visually, hence the results featured slightly higher counts than with visual scoring.

**Figure 7.**
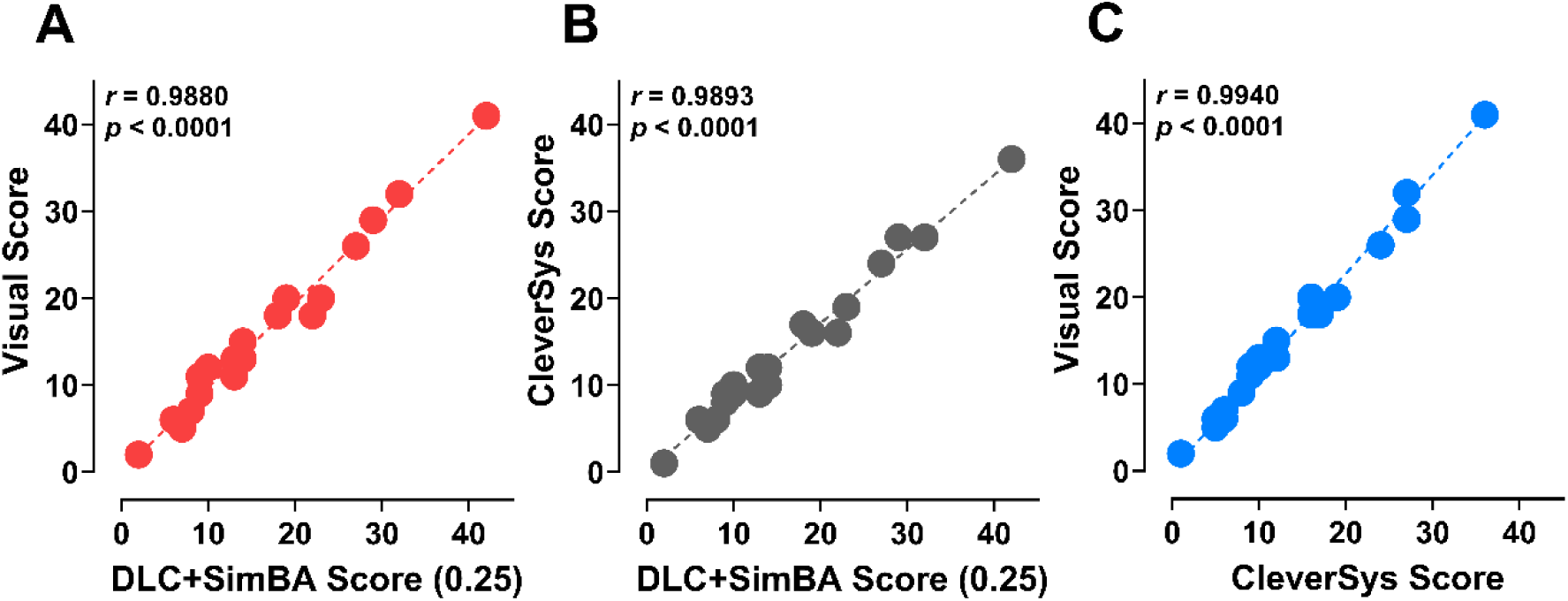
Correlations between HTR counts from 21 videos (bufotenine and WAY+bufotenine) across detection methods. (**A**) DLC+SimBA vs. visual scores, (**B**) commercial software vs. DLC+SimBA scores, and (**C**) visual scoring vs. commercial software scores.

Given the low HTR counts induced by bufotenine (max HTR counts <20 per 30 min session), we also sought to validate our approach for studying distinct psychedelic drug chemotypes producing variable levels of HTR counts. To accomplish this, additional experiments were conducted to measure HTR counts after administration of known HTR-active doses of psilocybin (1 mg/kg), DOI (1 mg/kg), and LSD (0.1 mg/kg) to C57BL/6J mice.*^75^* Time-course plots of the DLC+SimBA counts show that while HTRs induced by psilocybin and LSD peaked between 10 – 15 min post injection and returned to baseline by the end of the session, DOI HTR counts reached peak levels by 15 min and persisted throughout the session (**Figure 8A**). Total HTR counts were highest for the DOI treated mice (mean = 120), followed by LSD (mean = 52) and psilocybin (mean = 34) relative to vehicle controls (mean = 9; **Figure 8B**, **Table S7**), consistent with previous chemotype comparisons.

**Figure 8.**
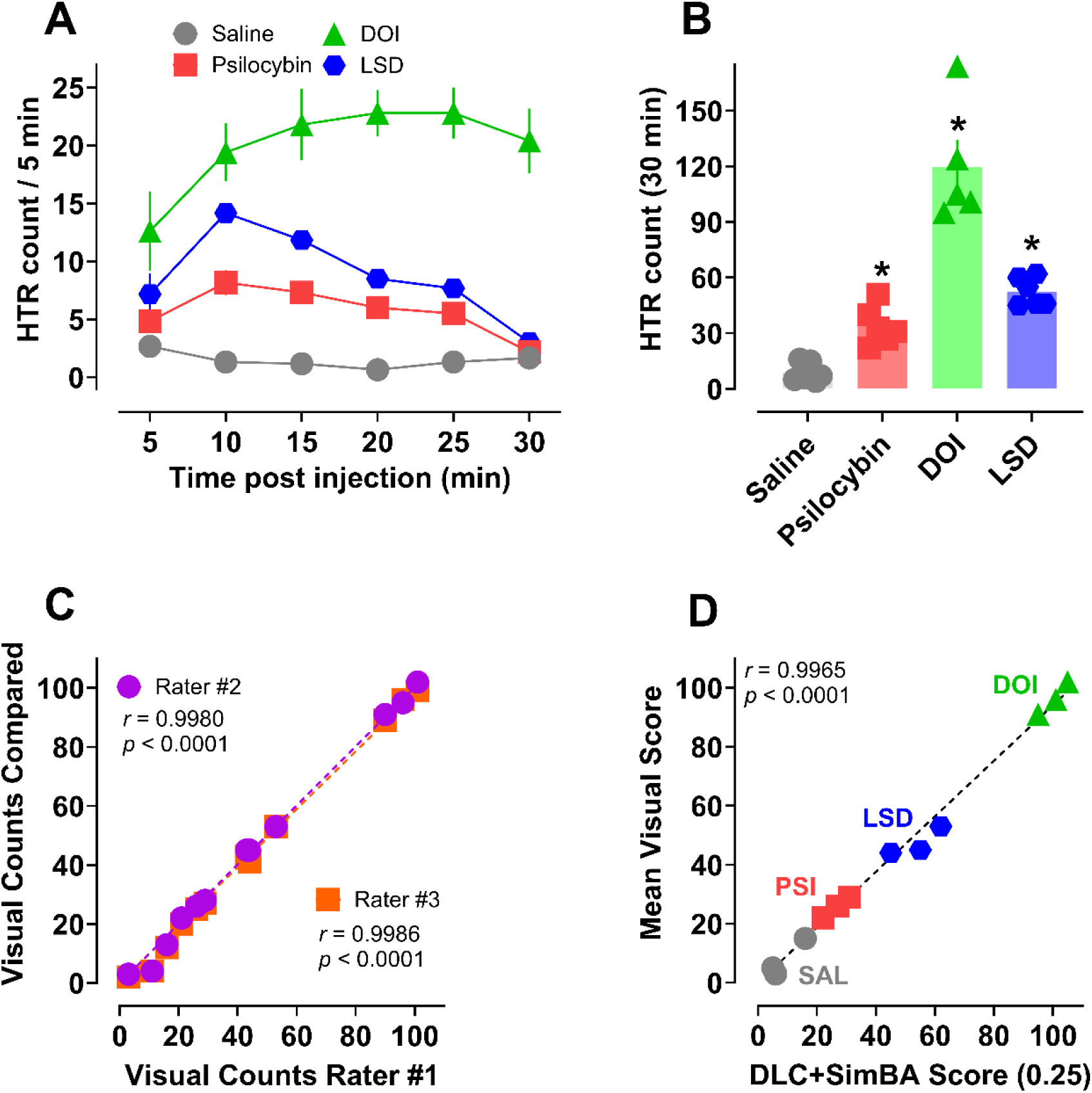
Additional HTR validation studies with psilocybin (1 mg/kg), DOI (1 mg/kg), and LSD (0.1 mg/kg). Time-course (**A**) and mean total counts (**B**) after s.c. administration of each drug. Correlations between three independent raters (**C**) and the mean visual score vs. DLC+SimBA HTR counts (**D**) from 12 videos. All values in **A** – **B** are mean ± SEM and represent *n* = 5 – 6 mice per data point. Asterisks represent statistical differences vs. saline vehicle controls (*p* < 0.05).

12 videos from the experiment (3 of each treatment group) were also visually scored by 3 independent raters blind to treatment conditions. The counts across raters #2 & 3 were strongly and significantly correlated with counts from rater #1, supporting similarity of the values across visual scores (**Figure 8C**). Furthermore, the mean visual score values for all 12 videos were also strongly and significantly correlated with counts from the DLC+SimBA approach (**Figure 8D**). It should be noted that the validation subset of 38 historical experimental videos used for evaluating each model combination also included variable doses of psychedelic chemotypes with established HTR activity. Based on the similarity of visual and DLC+SimBA scores, these results further validate the utility of our approach in characterizing psychedelic-like effects of 5-HT_2A_ agonists and importantly demonstrate that the approach can discriminate between compounds with variable HTR frequencies or total counts.

### 2.7 Advantages and limitations of using DLC + SimBA for HTR detection vs. other approaches

Here, we develop, validate, and deploy the deep learning-based pose estimation toolkit DLC and the behavioral classifier toolkit SimBA to demonstrate their utility as a rapid, accurate, and open-source approach to examine the behavioral effects of 5-HT_2A_ activity. The present approach has several key advantages over existing software-based approaches to HTR detection *^26^*, while overcoming potential limitations of magnetometer and existing software-based approaches.

Visual scoring of the HTR is time-consuming, typically requires multiple trained raters, and can exhibit inter-rater variability in scores.*^1, 74^* While quantification of HTRs has been reported using commercial software *^26^* and ML approaches *^28, 29^*, the present approach is the first to automate and validate this process with experimental videos in mice. Results from our best performing models suggest similar or improved performance metrics compared to the best performing DLC+SimBA model in the Cyrano and Popik *^29^* study in rats. However, the approaches are not directly comparable as the prior study focused on tracking the facial movements of rats from both sides of a cage, and rat wet-dog shakes have not been validated for predicting potency of psychedelic-like effects of drugs in humans. The only animal models that have been validated to predict potencies of psychedelics to induce subject effects in humans are drug discrimination in rats and the HTR paradigm in C57BL/6J mice.*^5^* Furthermore, rats do not respond the same across classes of psychedelics compared to C57BL/6J mice. For instance, the tryptamine psychedelics, psilocybin and DMT, produce measurable HTR in C57BL/6J mice *^5, 7, 76^*, but these drugs do not elicit appreciable wet-dog shakes in Sprague Dawley rats.*^77, 78^* Therefore, validation of the DLC+SimBA approach for studying the HTR in mice is an important advancement of available detection methods.

When comparing HTR detection methods, accuracy for the semi-automated software-based approach from Glatfelter et al. *^26^* was found to be 86 - 94% (depending on lighting conditions), which is slightly lower than values reported for magnetometer-based systems (96% - 99%) and the present approach (102%). Notably, the DLC+SimBA HTR scores were elevated due to a small number of true positive events missed by visual raters. Our validation testing suggests that our approach may detect the HTR more accurately than visual scoring in some cases, as some false positive HTR events were actually true positives missed by multiple trained visual raters. Regardless of minor differences in HTR counts, software- and magnetometer-based approaches generally agree on HTR potencies and relative maximal values observed across studies.*^5, 7, 38, 79^* However, accuracy measures alone may be insufficient for evaluating the robustness of an HTR detection method. A recent performance assessment of the modern magnetometer approach described originally by Halberstadt and Geyer *^1^* provided information regarding the number of HTR counts, false positives, and false negatives.*^25^* From the study results (see Table 1)*^25^*, the best calculated F_1_ score was ∼95%, comparable to the F_1_ score in our best performing model combination (96%) at 50% video resolution, 120 fps. Together, our results suggest that our DLC+SimBA HTR detection approach exhibits equivalent or greater performance compared to published assessments of other approaches.

Selecting an appropriate approach for HTR quantification may depend on the specific needs of investigators, given the limitations of the method described here and magnetometer-based HTR detection. For example, magnetometer-based approaches may be better suited for exclusively measuring the HTR where other drug effects are already well characterized. Magnetometer-based HTR counts are reported to be highly accurate and reliable across multiple laboratories. While our video-based approach is non-invasive, recently described magnetometer-based approaches using simple magnetic ear-tags are non-invasive, easy to build and attach, and require only brief anesthesia.*^23^* Furthermore, these systems can examine the dynamics of the head movements during individual HTR events.*^1, 5, 19, 21, 22^* Magnetometer systems can distinguish between subtypes of HTR events (e.g. short vs. long or low vs. high amplitude).*^22^* The present automated DLC+SimBA approach is limited in this regard, but it is possible that future iterations, perhaps with higher frame rate videos, may be able to accurately track the speed and dynamics of head movements during HTR events. Moreover, magnetometer-based approaches do not require any training data and provide nearly real-time data acquisition. Therefore, for investigators interested exclusively in HTR characterization and/or HTR dynamics, magnetometer-based methods remain the most robust and rapid approach.

In contrast, our approach, while automated, requires several hours for video analysis. Using the methods and equipment described here, it is possible to acquire data by the following morning using a model combination trained on additional HTR events at a lower resolution, if DLC analysis commences immediately following experiment completion. Importantly, the present approach is flexible and scalable beyond the specific methods described here. Using cloud-based computing resources, an institution’s high-performance computing cluster, or multiple graphical processing units (GPU), investigators can analyze experimental videos in parallel, significantly reducing data acquisition time to provide same-day results if desired. Future experiments should examine real-time pose estimation or other ML-based behavioral quantification approaches, such as with DLC-Live!, to further reduce data acquisition time.*^80^* Our approach lays the foundation for others to improve and expand these methods for even more rapid HTR detection and data acquisition comparable to magnetometer-based approaches.

Analysis of experimental videos using our approach may be particularly advantageous for characterization of novel compounds in the HTR paradigm with unknown potential for behavioral false positives. Since magnetometer-based systems do not record other behavioral effects of the drugs outside of the HTR, our approach also uniquely allows for simultaneous quantification of additional serotonergic behaviors, such as rearing, grooming, and ear scratching. Indeed, SimBA is designed for classification of multiple user-defined, complex animal behaviors.*^39^* Previous studies have successfully deployed DLC+SimBA to examine social behaviors in both rats and mice.*^81, 82^* Existing approaches are limited in detection of additional behaviors, while our approach allows investigators to interrogate experimental recordings for novel behavioral insights, which is especially relevant given the development of novel psychedelic drugs. It is well-established that indoleamine psychedelics are nonselective receptor agonists, and produce complex behavioral effects in rodents outside of the HTR that may warrant further investigation.*^6, 7, 64, 83–85^* Our approach provides the modularity to quantify and evaluate other behaviors of potential interest, by training new SimBA classifiers on top of existing HTR event annotations.

Another scenario where investigators may find our approach advantageous is for HTR quantification during behavioral assays such as fear conditioning, or testing the same animal later in a separate behavioral assay. Head-implanted or ear-attached magnets may interfere with behavioral assays due to magnets sticking to metal parts of testing chambers, or by potentially altering head movements. This distinction also highlights how our approach can be integrated into existing experimental setups and may be preferrable for investigators who already video record experimental sessions. Our results suggest that our approach can accurately quantify HTR even with lower resolution and frame rate video cameras, further highlighting the ease of integrating our approach.

The present validation studies also demonstrate that accurate detection of HTR can be achieved even with lower to moderate resolution and frame rate videos (50% resolution, 60 fps), suggesting that investigators can potentially use inexpensive video recording equipment. Excellent HTR detection performance was achieved using only 30 historical experimental videos to train DLC pose estimation models and 10 videos featuring 136 HTR events to train SimBA behavioral classifiers. The small amount of training data required for models with good performance highlights the accessibility of setting up the present approach. Additionally, both the DLC and SimBA toolkits are open-source with extensive documentation and software updates *^31, 39^*, and have been employed in diverse behavioral neuroscience paradigms.*^81, 82^* Future studies should examine using other open-source behavioral classification algorithms such as A-SoiD,*^86^* which requires significantly fewer human behavioral annotations due to the use of active learning, reducing training costs compared to annotating and training a SimBA behavioral classifier. Investigators should leverage the insights from our approach to guide the training, validation, and deployment of a DLC+SimBA model combination to detect HTR that is optimized for each laboratory.

To aid with deployment of this approach in other laboratories, several recommendations are provided below. First, as with the implementation of any novel assay, investigators should use reasonable controls, such as manually reviewing a small segment of experimental video to compare with visual HTR counts. Additionally, including a single dose positive control compound such as DOI or psilocybin (1 mg/kg) will allow comparison of counts across laboratories and scoring approaches. Second, while training a model and analyzing new videos with a DLC pose estimation model is computationally expensive and requires a mid-range GPU, using DLC with cloud-based computing resource services, such as Google Colab or Amazon Web Services, could bypass this problem. Investigators can easily train and deploy a pose estimation model for free or low cost, then run SimBA on their local computer. Third, to further enable “plug-and-play” of this approach, investigators may also consider implementing generalizable DLC models pre-trained to predict key points across animals (e.g., SuperAnimal models from the Mathis lab), which can readily track mice and be fine-tuned by investigators to accommodate variable experimental setups, further reducing the burden of annotating body parts for a new pose estimation model.*^87^*

## Conclusions

Our results show that using DLC+SimBA is a rapid, accurate, open-source, automated, and non-invasive approach for the assessment of HTRs in mice. This approach requires minimal training data and equipment, yet demonstrates excellent performance rivaling existing automated methods. Behavioral control experiments, robust validation testing, time cost comparisons, and post-hoc explainability metrics further support feasibility and reliability of the approach. To further assess the utility of this approach in psychedelic drug discovery, the HTR potency and efficacy of bufotenine was studied, revealing that 5-HT_1A_-mediated effects of the drug attenuate its effects on HTR in mice and that potent selective SERT release properties of bufotenine may be relevant to its overall pharmacological effects. Furthermore, the DLC+SimBA approach was successfully deployed to distinguish variable HTR frequencies induced by standard doses of psilocybin, DOI, and LSD. In summary, the present findings support the deployment of the DLC+SimBA HTR detection approach in mice and demonstrate key advantages over existing methods.

## Methods

### 3.1 Workflow overview and experimental design

For training and assessing our DLC and SimBA model combinations, 78 historical experimental videos were selected, primarily ranging from 12 to 17 minutes in length. Videos were randomly selected across animals, drug doses, session timepoints, cylinder alignments, lighting conditions, and experiment days to maximize training and testing set diversity for robust DLC and SimBA models. To avoid training-testing leakage, a common pitfall of ML-based approaches *^88^*, videos were randomly segregated into non-overlapping training and validation sets. For the final models, 30 videos were used for training the body part tracker using DLC (DLC training subset), 10 videos for training the behavioral classifier using SimBA (SimBA training subset), and 38 hold-out videos to assess the performance of the final DLC+SimBA model combinations for HTR detection by calculating the precision, recall, and F_1_ scores (training subset; **Figure 1B**). The validation subset videos included 18 experimental videos from Glatfelter et al. *^26^*, and 20 additional random historical experimental videos across various psychedelic drugs (i.e. psilocybin, LSD, DOI, DOBU, 2,5-DMA, DOPR), doses (0.001 – 30 mg/kg), lighting conditions (standard room overhead and enhanced lighting, i.e. using highest illumination setting on GoPro Zeus Mini magnetic swivel clip accessory lights placed above the open field arenas as described), and animals (male and female), for a total of 38 experimental videos (**Figure 1B**).*^75, 89^* Each of the 9 behavioral classifiers were assessed at various discrimination thresholds, from 0% to 50%. The videos used for training and initial validation (78 videos) had 67% male mice, whereas deployment videos had equal numbers of males and females. The 18 validation videos from Glatfelter et al. (2022) used only male subjects. For the remaining videos (60 videos), 54% of the mice were male.

Briefly, all drug experimental videos used the following parameters, as described in detail by Glatfelter et al. *^26^*. Hero Black 7 GoPro cameras (GoPro Inc.) were used to record 120 fps videos at 1280 x 960 resolution. For the experimental sessions, mice received drug or vehicle injections (0.01 mL/g body weight) and were placed into cylindrical acrylic arenas (7.5 in diameter) housed inside of TruScan mouse locomotor boxes (Coulbourn Instruments). The arenas had transparent floor panels with white bench paper underneath to provide a light background for contrast. GoPro cameras were mounted ∼ 10 inches above the arena floor, and experimental test sessions were recorded for 30 min post injection. All experiments occurred during the light phase of the light–dark cycle between 0800 and 1700 h local time (lights on at 0600). Subjects were randomized to treatment conditions and were repeat tested once per 1–2 weeks to avoid tolerance to the effects of each drug on HTR. After videos of each experiment were recorded, the video files were transferred to an external hard drive for storage until subsequent training and analysis as described below. All experiments were approved by the NIDA IRP Animal Care and Use Committee under protocol 23-OSD-37.

### 3.2 Video pre-processing

To assess combined DLC+SimBA model performances across video resolutions and frame rates, all 78 videos were batch preprocessed using FFmpeg, a free, open-source software tool for handling video files (https://www.ffmpeg.org/). The video files were downscaled from 1280 x 960 pixels (100% resolution), 120 fps to 906 x 680 pixels (50% resolution), 640 x 480 pixels (25% resolution), and 454 x 340 pixels (12.5% resolution) at 120, 60, and 30 fps. In summary, for training and assessing the DLC and SimBA models, there were 9 sets of 78 videos, representing each resolution and frame rate.

### 3.3 Mouse pose estimation models

For body part tracking, DLC (version 2.3) [(Mathis et al. 2018), (Nath et al. 2019)] was used for training a model at each video resolution (50%, 25%, 12.5%). Specifically, extracted frames taken from 30 videos (at each resolution) and 16 animals (then 95% was used for training) were labeled. The number of total extracted frames was 600 frames for each DLC model. Thirteen body points were consistently labeled, including nose, left eye, right eye, left ear, right ear, top of the skull, three points along the spine, the tail base, and three points along the tail (**Figure 1A**). A ResNet-50-based neural network *^90, 91^* was used with default parameters (batch size of 8) for 500,000 training iterations, when loss typically plateaus.*^30^* Each model was validated with 3 shuffles. The train and test errors for each model can be found in **Table S1**, as well as comparisons of each models’ analysis times in **Table S2**. Prior to behavioral classifier training using SimBA, two internal controls videos of an empty cylinder arena and a historical experimental video featuring an immobile mouse were analyzed to further validate each DLC model.

### 3.4 Behavioral classifiers

Behavioral classifiers were generated with 13 user-defined body parts in SimBA for each video resolution and frame rate, for a total of 9 behavioral classifiers. Since the same set of historical experimental videos were used, the HTR event annotations were copied across resolutions. For example, the behavioral classifier trained at 50% resolution, 30 fps had identical behavior annotations as the behavioral classifier trained at 25% resolution, 30 fps. The approximate width of the TruScan chamber was used to define pixels per mm during the Video Settings step. The outlier correction step was skipped. Features derived from mouse pose estimation data (32 features) were extracted using default settings.

Classifier training videos were carefully annotated for the HTR using the default SimBA behavioral annotator GUI. A total of 1,212,960 frames (at 120 fps), 606,480 frames (at 60 fps), and 303,240 frames (at 30 fps) from 10 random historical experimental videos at each resolution (2.8 h of video recording) were annotated for HTR events. Each HTR was annotated from the initial movement of the mouse’s ears until the head returned to its initial resting state. Frequencies of annotated HTR behavior in the training subset were 0.13% at 30 fps, 0.1% at 60 fps, and 0.09% at 120 fps. To further confirm model performance was not dependent on a specific labeler, the same classifier training videos were also carefully annotated for HTR by another experimenter using methods described above. Random forest models were generated in SimBA with the default hyperparameters (algorithm = RF, random forest estimators = 2000, max features = sqrt, criterion = gini, test size = 0.2, train-test split type = frames, minimum sample leaf = 1) and included optional model evaluation settings to generate a classification report, precision recall curves, feature importance bar graph, and SHAP score calculations for 100 frames present and 100 frames absent.

### 3.5 Post-hoc explainability metrics for behavioral classifiers

Feature importance values were calculated for each behavioral classifier in SimBA. SHAP analyses were run on 100 random frames with HTR behavior present and 100 random frames with HTR behavior absent for each model. Feature importance values are provided in **Figure 4**. SHAP value correlations can be provided upon reasonable request.

### 3.6 Mouse experiments

Male and female C57BL/6J mice were purchased at 8 weeks of age from a commercial vendor (The Jackson Laboratory #000664) and were initially group housed (4/cage) at the NIDA IRP animal facilities in Baltimore, MD for 1 – 2 weeks prior to experiments. After acclimation to the housing facility, each mouse was subjected to brief isoflurane anesthesia and a TP500 transponder (Avidity Science) was implanted subcutaneously on the upper back as previously described.*^7, 92^* The transponder served as an animal identifier and allowed remote, noninvasive reading of body temperature using a handheld reader (Avidity Science, model DAS-8027 IUS). After implantation, mice were single housed to prevent removal of the transponder by cage mates.

### 3.7 Behavioral control experimental videos

Previously recorded experimental videos of d-amphetamine-induced locomotion and stereotypy and SKF-38393-induced grooming from Glatfelter et al. *^26^* were analyzed using the best performing DLC and SimBA model combination. For detailed methods for drug-induced behavioral control experiments, see Glatfelter et al. *^26^*.

### 3.8 Bufotenine dose-response and validation experiments

Prior to each experiment, mouse body weight and temperature were recorded. Mice were then placed into testing chambers for acclimation. In dose–response studies, after a brief 5 min acclimation, mouse body temperature was recorded for baseline measurement, mice received a s.c. injection of bufotenine (Cayman Chemical #11803 “neat” preparation) or saline vehicle, and animals were returned to the testing arena for 30 min. Similarly, validation experiments entailed s.c. administration of DOI HCl (1 mg/kg, Cayman Chemical #13885), (+)-LSD tartrate (2:1) (0.1 mg/kg, NIDA drug supply program), or psilocybin (1 mg/kg, NIDA drug supply program) under the same conditions. During the sessions, locomotor activity was monitored via photobeam tracking of movements in the horizontal plane to yield distance traveled in centimeters. Video recordings used GoPro Hero Black 7 cameras (1280 x 960, 120 fps). Post-treatment body temperature values were also recorded. In antagonist reversal experiments, mice received a s.c. injection of either WAY100635 maleate (Cayman Chemical #14599) or saline vehicle 10 mins prior to bufotenine and were returned to the testing chamber for 30 mins. During this period, locomotor activity was monitored to examine the potential effects of antagonist treatment on general behavior or movement. All drug doses represent the weight of the salt form used.

### 3.9 Computer hardware and software for machine learning models

All models were trained on a Dell Precision Workstation T5820 with Intel Xeon Processor W-2223 (3.6GHz), 32 GB RAM, Windows 11 Pro operating system, and a NVIDIA RTX A100 GPU (video card). Videos were downscaled using FFmpeg. Pose estimation models were trained using DLC version 2.3.10. Behavioral classifiers were generated using SimBA version 2.2.4.

### 3.10 Data Analysis and Statistics

Data for the acute effects of bufotenine on HTR were fit with biphasic dose-response nonlinear regression fits for visualization. Data for acute effects of bufotenine on body temperature change and motor activity were fit with four parameter nonlinear regression fits (IC_50_ > 0) except the motor activity for WAY pretreatment. Potency values for HTR were determined using four parameter nonlinear regression fits of ascending limb doses of bufotenine+WAY (EC_50_ > 0). Comparisons of dose-response data for acute drug effects (HTR, body temperature change, motor activity) were conducted via one-way (equal variance) or Welch’s (unequal variance) ANOVA followed by Dunnett’s or Dunnett’s T3 post hoc analyses comparing all groups to vehicle. Predetermined dose by dose comparisons were made between no pretreatment and WAY pretreatment groups using a two-way ANOVA (dose x treatment) with Sidak’s post hoc test. Pearson correlation analyses (two tailed) were conducted to assess relationships between HTR counts across scoring methods. Alpha was set at 0.05 for all analyses. Three parameter nonlinear regression fits were used to determine best fit values of interest (IC_50_, EC_50_, E_max_) in monoamine transporter assays.

### 3.11 In Vitro Transporter Assays

Monoamine transporter assays were conducted as previously described.*^53, 93–95^* Briefly, Sprague-Dawley rats were euthanized via CO_2_ narcosis and brain tissue was collected. Caudate tissue was used for dopamine transporter assays, while whole brain minus caudate and cerebellum was used for norepinephrine transporter and SERT assays. Synaptosomes were prepared using standard methods and utilized for both uptake inhibition and release assays. [^3^H]5-HT, [^3^H]dopamine, and [^3^H]norepinephrine were used for uptake inhibition assays at their respective transporters. For release assays, [^3^H]5-HT was used as the radiolabeled substrate for SERT, while [^3^H]MPP+ was used as the radiolabeled substrate for dopamine transporters and norepinephrine transporters.

## Author Contributions

Experimental design – A.D.M, G.C.G., M.H.B.

Scoring and validation – A.D.M, D.W., N.R.G., G.C.G.

Mouse experiments – G.C.G., A.D.M., N.R.G.

Data analyses – G.C.G., A.D.M., F. P.

Initial draft – A.D.M., G.C.G.

Editing and proofreading – G.C.G., A.D.M., M.H.B., F.P., N.R.G, D.W.

## Abbreviations

DLC: DeepLabCut
DOI: 2,5-dimethoxy-4-iodoamphetamine
DOBU: 2,5-dimethoxy-4-butylamphetamine
DOPR: 2,5-dimethoxy-4-propylamphetamine
2,5-DMA: 2,5-dimethoxyamphetamine
LSD: lysergic acid diethylamide
Fps: frames per second
HTR: Head-twitch response
ML: Machine learning
SimBA: Simple Behavioral Analysis
5-HT: Serotonin
5-HT_1A_: Serotonin 1A receptors
5-HT_2A_: Serotonin 2A receptor
SERT: Serotonin transporter
5-HO-DMT: 5-hydroxy-N,N-dimethyltryptamine
5-MeO-DMT: 5-methoxy-*N*,*N*-dimethyltryptamine
DMT: *N*,*N*-dimethyltryptamine
WAY: WAY100635

## Supporting Information

Final snapshot table for the DLC model at each video resolution; DLC inference time and FFmpeg downscaling time cost table; DLC pose estimation model time costs for 50% resolution (906 x 680); SimBA behavioral classifier model training time in minutes for 50% resolution (906 x 680), 120 fps model; FFmpeg video downscaling time, DLC inference time, and SimBA video analysis for HTR quantification times in minutes for 50% resolution; 120 fps model combination at 0.25 discrimination threshold; 120 fps videos heatmap plots of accuracy-discrimination threshold curve summary data; accuracy-discrimination threshold curves from a second experimenter; heatmap plots of accuracy-discrimination threshold curve summary data from a second experimenter; time-course plots for acute effects of bufotenine on the HTR; time-course plots for acute effects of bufotenine on motor activity; summary statistics table for acute effects of bufotenine; summary statistics table for further validation studies; heatmap plots illustrating monoamine transporter activity of bufotenine.

## Acknowledgements

This work was supported by NIDA Intramural Research Program grant number DA-000522-16 (M.H.B.). We would like to thank the Mathis Laboratory and Golden Laboratory for their open-source toolkits, DLC and Simple Behavioral Analysis respectively. We would like to especially thank Simon Nilsson of the Golden Laboratory for their insights on the community Github forums. The structure in **Figure 6A** was created using ChemDraw (v 21.0.0).

---

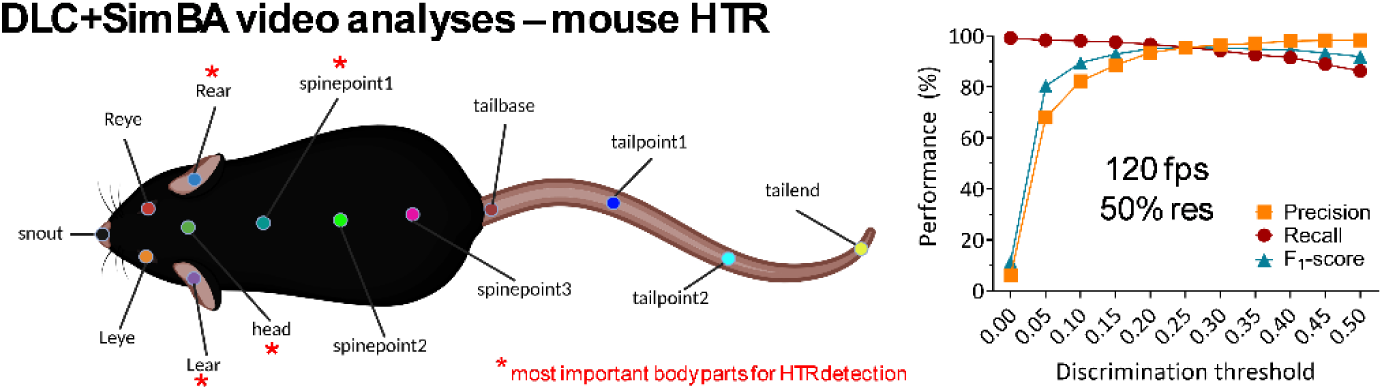

**Synopsis:** All body points used to train pose-estimation DLC models with key points used to detect head twitch responses noted with red asterisks. Performance metrics across discrimination thresholds for the DLC+SimBA model combination to analyze 50% resolution 120 fps videos.

## References

(1) Halberstadt, A. L., and Geyer, M. A. (2013) Characterization of the head-twitch response induced by hallucinogens in mice: detection of the behavior based on the dynamics of head movement, Psychopharmacology (Berl) 227, 727–739, DOI: 10.1007/s00213-013-3006-z.

(2) González-Maeso, J., Weisstaub, N. V., Zhou, M., Chan, P., Ivic, L., Ang, R., Lira, A., Bradley-Moore, M., Ge, Y., Zhou, Q., Sealfon, S. C., and Gingrich, J. A. (2007) Hallucinogens Recruit Specific Cortical 5-HT2A Receptor-Mediated Signaling Pathways to Affect Behavior, Neuron 53, 439–452, DOI: 10.1016/j.neuron.2007.01.008.

(3) Canal, C. E., and Morgan, D. (2012) Head-twitch response in rodents induced by the hallucinogen 2,5-dimethoxy-4-iodoamphetamine: a comprehensive history, a re-evaluation of mechanisms, and its utility as a model, Drug Test Anal 4, 556–576, DOI: 10.1002/dta.1333.

(4) Nichols, D. E. (2016) Psychedelics, Pharmacological Reviews 68, 264–355, DOI: 10.1124/pr.115.011478.

(5) Halberstadt, A. L., Chatha, M., Klein, A. K., Wallach, J., and Brandt, S. D. (2020) Correlation between the potency of hallucinogens in the mouse head-twitch response assay and their behavioral and subjective effects in other species, Neuropharmacology 167, 107933.

(6) Glatfelter, G. C., Clark, A. A., Cavalco, N. G., Landavazo, A., Partilla, J. S., Naeem, M., Golen, J. A., Chadeayne, A. R., Manke, D. R., Blough, B. E., McCorvy, J. D., and Baumann, M. H. (2024) Serotonin 1A Receptors Modulate Serotonin 2A Receptor-Mediated Behavioral Effects of 5-Methoxy-N,N-dimethyltryptamine Analogs in Mice, ACS Chemical Neuroscience, DOI: 10.1021/acschemneuro.4c00513.

(7) Glatfelter, G. C., Pottie, E., Partilla, J. S., Sherwood, A. M., Kaylo, K., Pham, D. N. K., Naeem, M., Sammeta, V. R., DeBoer, S., Golen, J. A., Hulley, E. B., Stove, C. P., Chadeayne, A. R., Manke, D. R., and Baumann, M. H. (2022) Structure-Activity Relationships for Psilocybin, Baeocystin, Aeruginascin, and Related Analogues to Produce Pharmacological Effects in Mice, ACS Pharmacol Transl Sci 5, 1181–1196, DOI: 10.1021/acsptsci.2c00177.

(8) Varty, G. B., Canal, C. E., Mueller, T. A., Hartsel, J. A., Tyagi, R., Avery, K., Morgan, M. E., Reichelt, A. C., Pathare, P., Stang, E., Palfreyman, M. G., and Nivorozhkin, A. (2024) Synthesis and Structure–Activity Relationships of 2,5-Dimethoxy-4-Substituted Phenethylamines and the Discovery of CYB210010: A Potent, Orally Bioavailable and Long-Acting Serotonin 5-HT2 Receptor Agonist, Journal of Medicinal Chemistry 67, 6144–6188, DOI: 10.1021/acs.jmedchem.3c01961.

(9) Nichols, D. E., Sassano, M. F., Halberstadt, A. L., Klein, L. M., Brandt, S. D., Elliott, S. P., and Fiedler, W. J. (2015) N-Benzyl-5-methoxytryptamines as Potent Serotonin 5-HT2 Receptor Family Agonists and Comparison with a Series of Phenethylamine Analogues, ACS Chemical Neuroscience 6, 1165–1175, DOI: 10.1021/cn500292d.

(10) Halberstadt, A. L., and Geyer, M. A. (2014) Effects of the hallucinogen 2, 5-dimethoxy-4-iodophenethylamine (2C-I) and superpotent N-benzyl derivatives on the head twitch response, Neuropharmacology 77, 200–207.

(11) Sherwood, A. M., Burkhartzmeyer, E. K., Williamson, S. E., Baumann, M. H., and Glatfelter, G. C. (2024) Psychedelic-like Activity of Norpsilocin Analogues, ACS Chemical Neuroscience 15, 315–327, DOI: 10.1021/acschemneuro.3c00610.

(12) Fantegrossi, W. E., Simoneau, J., Cohen, M. S., Zimmerman, S. M., Henson, C. M., Rice, K. C., and Woods, J. H. (2010) Interaction of 5-HT2A and 5-HT2C receptors in R(-)-2,5-dimethoxy-4-iodoamphetamine-elicited head twitch behavior in mice, J Pharmacol Exp Ther 335, 728–734, DOI: 10.1124/jpet.110.172247.

(13) Cameron, L. P., Tombari, R. J., Lu, J., Pell, A. J., Hurley, Z. Q., Ehinger, Y., Vargas, M. V., McCarroll, M. N., Taylor, J. C., Myers-Turnbull, D., Liu, T., Yaghoobi, B., Laskowski, L. J., Anderson, E. I., Zhang, G., Viswanathan, J., Brown, B. M., Tjia, M., Dunlap, L. E., Rabow, Z. T., Fiehn, O., Wulff, H., McCorvy, J. D., Lein, P. J., Kokel, D., Ron, D., Peters, J., Zuo, Y., and Olson, D. E. (2021) A non-hallucinogenic psychedelic analogue with therapeutic potential, Nature 589, 474–479, DOI: 10.1038/s41586-020-3008-z.

(14) Darmani, N. A., Shaddy, J., and Gerdes, C. F. (1996) Differential Ontogenesis of Three DOI-Induced Behaviors in Mice, Physiology & Behavior 60, 1495–1500, DOI: 10.1016/S0031-9384(96)00323-X.

(15) Erkizia-Santamaría, I., Alles-Pascual, R., Horrillo, I., Meana, J. J., and Ortega, J. E. (2022) Serotonin 5-HT2A, 5-HT2c and 5-HT1A receptor involvement in the acute effects of psilocybin in mice. In vitro pharmacological profile and modulation of thermoregulation and head-twich response, Biomedicine & Pharmacotherapy 154, 113612, DOI: 10.1016/j.biopha.2022.113612.

(16) Canal, C. E., Olaghere da Silva, U. B., Gresch, P. J., Watt, E. E., Sanders-Bush, E., and Airey, D. C. (2010) The serotonin 2C receptor potently modulates the head-twitch response in mice induced by a phenethylamine hallucinogen, Psychopharmacology (Berl) 209, 163–174, DOI: 10.1007/s00213-010-1784-0.

(17) Holloway, T., Moreno, J. L., and González-Maeso, J. (2016) HSV-Mediated Transgene Expression of Chimeric Constructs to Study Behavioral Function of GPCR Heteromers in Mice. Journal of visualized experiments : JoVE DOI: 10.3791/53717.

(18) U.S. Department of Health and Human Services: Office of Disease Prevention and Health Promotion--Healthy People 2020, Nasnewsletter 15, 3.

(19) de la Fuente Revenga, M., Shin, J. M., Vohra, H. Z., Hideshima, K. S., Schneck, M., Poklis, J. L., and González-Maeso, J. (2019) Fully automated head-twitch detection system for the study of 5-HT(2A) receptor pharmacology in vivo, Sci Rep 9, 14247, DOI: 10.1038/s41598-019-49913-4.

(20) Jaster, A. M., and González-Maeso, J. (2023) Automated Detection of Psychedelic-Induced Head-Twitch Response in Mice. In Schizophrenia: Methods and Protocols (Urigüen, L., and Díez-Alarcia, R., Eds.), pp 65–76, Springer US, New York, NY.

(21) de la Fuente Revenga, M., Vohra, H. Z., and González-Maeso, J. (2020) Automated quantification of head-twitch response in mice via ear tag reporter coupled with biphasic detection, Journal of Neuroscience Methods 334, 108595, DOI: 10.1016/j.jneumeth.2020.108595.

(22) Halberstadt, A. L. (2020) Automated detection of the head-twitch response using wavelet scalograms and a deep convolutional neural network, Scientific reports 10, 8344–8344, DOI: 10.1038/s41598-020-65264-x.

(23) Jefferson, S. J., Gregg, I., Dibbs, M., Liao, C., Wu, H., Davoudian, P. A., Woodburn, S. C., Wehrle, P. H., Sprouse, J. S., Sherwood, A. M., Kaye, A. P., Pittenger, C., and Kwan, A. C. (2023) 5-MeO-DMT modifies innate behaviors and promotes structural neural plasticity in mice, Neuropsychopharmacology 48, 1257–1266, DOI: 10.1038/s41386-023-01572-w.

(24) Siegel, R. K., Lee, M. A., and Jarvik, M. E. (1972) A device for analyzing drug-induced responses in freely moving mice, Journal of the experimental analysis of behavior 18, 415–418, DOI: 10.1901/jeab.1972.18-415.

(25) Nakamura, M., Hojo, M., Kawai, A., Ikushima, K., Nagasawa, A., Takahashi, H., Makino, K., Suzuki, T., Suzuki, J., and Inomata, A. (2023) An application of the magnetometer detection system to Crl:CD1 (ICR) mice for head twitch response induced by hallucinogenic 5-HT 2A agonists, Fundamental Toxicological Sciences 10, 189–197, DOI: 10.2131/fts.10.189.

(26) Glatfelter, G. C., Chojnacki, M. R., McGriff, S. A., Wang, T., and Baumann, M. H. (2022) Automated Computer Software Assessment of 5-Hydroxytryptamine 2A Receptor-Mediated Head Twitch Responses from Video Recordings of Mice, ACS Pharmacology & Translational Science, DOI: 10.1021/acsptsci.1c00237.

(27) Sturman, O., von Ziegler, L., Schläppi, C., Akyol, F., Privitera, M., Slominski, D., Grimm, C., Thieren, L., Zerbi, V., Grewe, B., and Bohacek, J. (2020) Deep learning-based behavioral analysis reaches human accuracy and is capable of outperforming commercial solutions, Neuropsychopharmacology 45, 1942–1952, DOI: 10.1038/s41386-020-0776-y.

(28) Contreras, A., Khumnark, M., Hines, R. M., and Hines, D. J. (2021) Behavioral arrest and a characteristic slow waveform are hallmark responses to selective 5-HT2A receptor activation, Scientific Reports 11, 1925, DOI: 10.1038/s41598-021-81552-6.

(29) Cyrano, E., and Popik, P. (2025) Assessing the effects of 5-HT2A and 5-HT5A receptor antagonists on DOI-induced head-twitch response in male rats using marker-less deep learning algorithms, Pharmacological Reports 77, 135–144, DOI: 10.1007/s43440-024-00679-1.

(30) Mathis, A., Mamidanna, P., Cury, K. M., Abe, T., Murthy, V. N., Mathis, M. W., and Bethge, M. (2018) DeepLabCut: markerless pose estimation of user-defined body parts with deep learning, Nature Neuroscience 21, 1281–1289, DOI: 10.1038/s41593-018-0209-y.

(31) Nath, T., Mathis, A., Chen, A. C., Patel, A., Bethge, M., and Mathis, M. W. (2019) Using DeepLabCut for 3D markerless pose estimation across species and behaviors, Nature Protocols 14, 2152–2176, DOI: 10.1038/s41596-019-0176-0.

(32) Shulgin, A. T., and Shulgin, A. (1997) TiHKAL: The Continuation, Transform Press.

(33) Ott, J. (2001) Pharmañopo—Psychonautics: Human Intranasal, Sublingual, Intrarectal, Pulmonary and Oral Pharmacology of Bufotenine, Journal of Psychoactive Drugs 33, 273–281, DOI: 10.1080/02791072.2001.10400574.

(34) Fabing, H. D., and Hawkins, J. R. (1956) Intravenous Bufotenine Injection in the Human Being, Science 123, 886–887, DOI: doi:10.1126/science.123.3203.886.

(35) McLeod, W. R., and Sitaram, B. R. (1985) Bufotenine reconsidered, Acta Psychiatrica Scandinavica 72, 447–450, DOI: 10.1111/j.1600-0447.1985.tb02638.x.

(36) McBride, M. C. (2000) Bufotenine: Toward an Understanding of Possible Psychoactive Mechanisms, Journal of Psychoactive Drugs 32, 321–331, DOI: 10.1080/02791072.2000.10400456.

(37) Glatfelter, G. C., Naeem, M., Pham, D. N. K., Golen, J. A., Chadeayne, A. R., Manke, D. R., and Baumann, M. H. (2023) Receptor Binding Profiles for Tryptamine Psychedelics and Effects of 4-Propionoxy-N,N-dimethyltryptamine in Mice, ACS Pharmacology & Translational Science, DOI: 10.1021/acsptsci.2c00222.

(38) Glatfelter, G. C., Pottie, E., Partilla, J. S., Stove, C. P., and Baumann, M. H. (2024) Comparative Pharmacological Effects of Lisuride and Lysergic Acid Diethylamide Revisited, ACS Pharmacology & Translational Science 7, 641–653, DOI: 10.1021/acsptsci.3c00192.

(39) Goodwin, N. L., Choong, J. J., Hwang, S., Pitts, K., Bloom, L., Islam, A., Zhang, Y. Y., Szelenyi, E. R., Tong, X., Newman, E. L., Miczek, K., Wright, H. R., McLaughlin, R. J., Norville, Z. C., Eshel, N., Heshmati, M., Nilsson, S. R. O., and Golden, S. A. (2024) Simple Behavioral Analysis (SimBA) as a platform for explainable machine learning in behavioral neuroscience, Nature Neuroscience 27, 1411–1424, DOI: 10.1038/s41593-024-01649-9.

(40) Breiman, L., Friedman, J., Olshen, R. A., and Stone, C. J. (1984) Classification and Regression Trees, 1st ed., Chapman and Hall/CRC DOI: 10.1201/9781315139470.

(41) Goodwin, N. L., Nilsson, S. R. O., Choong, J. J., and Golden, S. A. (2022) Toward the explainability, transparency, and universality of machine learning for behavioral classification in neuroscience, Current Opinion in Neurobiology 73, 102544, DOI: 10.1016/j.conb.2022.102544.

(42) Zhang, M., Yang, Y., Yang, Z., Wen, X., Zhang, C., Xiao, P., Wang, Y., Sun, J., Wang, H., and Wang, X. (2025) Structural insights into tryptamine psychedelics: The role of hydroxyl indole ring site in 5-HT2A receptor activation and psychedelic-like activity, European Journal of Medicinal Chemistry 281, 117049, DOI: 10.1016/j.ejmech.2024.117049.

(43) Puigseslloses, P., Nadal-Gratacós, N., Ketsela, G., Weiss, N., Berzosa, X., Estrada-Tejedor, R., Islam, M. N., Holy, M., Niello, M., Pubill, D., Camarasa, J., Escubedo, E., Sitte, H. H., and López-Arnau, R. (2024) Structure-activity relationships of serotonergic 5-MeO-DMT derivatives: insights into psychoactive and thermoregulatory properties, Molecular Psychiatry, DOI: 10.1038/s41380-024-02506-8.

(44) Lyttle, T., David, G., and and Gartz, J. (1996) Bufo Toads and Bufotenine: Fact and Fiction Surrounding an Alleged Psychedelic, Journal of Psychoactive Drugs 28, 267–290, DOI: 10.1080/02791072.1996.10472488.

(45) Shulgin, A. T. (1981) Bufotenine, Journal of Psychoactive Drugs 13, 389–389, DOI: 10.1080/02791072.1981.10471899.

(46) Turner, W. J., and Merlis, S. (1959) Effect of Some Indolealkylamines on Man, A.M.A. Archives of Neurology & Psychiatry 81, 121–129, DOI: 10.1001/archneurpsyc.1959.02340130141020.

(47) Chilton, W. S., Jeremy, B., and and Jensen, R. E. (1979) Psilocin, Bufotenine and Serotonin: Historical and Biosynthetic Observations, Journal of Psychedelic Drugs 11, 61–69, DOI: 10.1080/02791072.1979.10472093.

(48) Delices, M., Muller, J. d. A. I., Arunachalam, K., and Martins, D. T. d. O. (2023) Anadenanthera colubrina (Vell) Brenan: Ethnobotanical, phytochemical, pharmacological and toxicological aspects, Journal of Ethnopharmacology 300, 115745, DOI: 10.1016/j.jep.2022.115745.

(49) Moretti, C., Gaillard, Y., Grenand, P., Bévalot, F., and Prévosto, J.-M. (2006) Identification of 5-hydroxy-tryptamine (bufotenine) in takini (Brosimum acutifolium Huber subsp. acutifolium C.C. Berg, Moraceae), a shamanic potion used in the Guiana Plateau, Journal of Ethnopharmacology 106, 198–202, DOI: 10.1016/j.jep.2005.12.022.

(50) Barry, T. L., Petzinger, G., and Zito, S. W. (1996) GC/MS comparison of the West Indian aphrodisiac "Love Stone" to the Chinese medication "chan su": bufotenine and related bufadienolides, J Forensic Sci 41, 1068–1073.

(51) Chamakura, R. P. (1994) Bufotenine - A Hallucinogen in Ancient Snuff Powders of South America and a Drug of Abuse on the Streets of New York City, Forensic Sci Rev 6, 1–18.

(52) Barker, S. A., McIlhenny, E. H., and Strassman, R. (2012) A critical review of reports of endogenous psychedelic N, N-dimethyltryptamines in humans: 1955–2010, Drug Testing and Analysis 4, 617–635, DOI: 10.1002/dta.422.

(53) Blough, B. E., Landavazo, A., Decker, A. M., Partilla, J. S., Baumann, M. H., and Rothman, R. B. (2014) Interaction of psychoactive tryptamines with biogenic amine transporters and serotonin receptor subtypes, Psychopharmacology 231, 4135–4144, DOI: 10.1007/s00213-014-3557-7.

(54) Roth, B. L., Choudhary, M. S., Khan, N., and Uluer, A. Z. (1997) High-affinity agonist binding is not sufficient for agonist efficacy at 5-hydroxytryptamine2A receptors: evidence in favor of a modified ternary complex model, J Pharmacol Exp Ther 280, 576–583.

(55) Choudhary, M. S., Craigo, S., and Roth, B. L. (1993) A single point mutation (Phe340-->Leu340) of a conserved phenylalanine abolishes 4-[125I]iodo-(2,5-dimethoxy)phenylisopropylamine and [3H]mesulergine but not [3H]ketanserin binding to 5-hydroxytryptamine2 receptors, Mol Pharmacol 43, 755–761.

(56) Pauwels, P. J., Van Gompel, P., and Leysen, J. E. (1993) Activity of serotonin (5-HT) receptor agonists, partial agonists and antagonists at cloned human 5-HT1a receptors that are negatively coupled to adenylate cyclase in permanently transfected hela cells, Biochemical Pharmacology 45, 375–383, DOI: 10.1016/0006-2952(93)90073-6.

(57) Williams, G. M., Smith, D. L., and Smith, D. J. (1992) 5-HT3 receptors are not involved in the modulation of the K+-evoked release of [3H]5-HT from spinal cord synaptosomes of rat, Neuropharmacology 31, 725–733, DOI: 10.1016/0028-3908(92)90033-L.

(58) Peroutka, S. J. (1985) Selective labeling of 5-HT1A and 5-HT1B binding sites in bovine brain, Brain Research 344, 167–171, DOI: 10.1016/0006-8993(85)91204-1.

(59) Kozell, L. B., Eshleman, A. J., Swanson, T. L., Bloom, S. H., Wolfrum, K. M., Schmachtenberg, J. L., Olson, R. J., Janowsky, A. J., and Abbas, A. I. (2023) Pharmacologic activity of substituted tryptamines at 5-HT(2A)R, 5-HT(2C)R, 5-HT(1A)R, and SERT, J Pharmacol Exp Ther, DOI: 10.1124/jpet.122.001454.

(60) Cozzi, N. V., Gopalakrishnan, A., Anderson, L. L., Feih, J. T., Shulgin, A. T., Daley, P. F., and Ruoho, A. E. (2009) Dimethyltryptamine and other hallucinogenic tryptamines exhibit substrate behavior at the serotonin uptake transporter and the vesicle monoamine transporter, Journal of Neural Transmission 116, 1591–1599, DOI: 10.1007/s00702-009-0308-8.

(61) Becker, A. M., Klaiber, A., Holze, F., Istampoulouoglou, I., Duthaler, U., Varghese, N., Eckert, A., and Liechti, M. E. (2023) Ketanserin Reverses the Acute Response to LSD in a Randomized, Double-Blind, Placebo-Controlled, Crossover Study in Healthy Participants, Int J Neuropsychopharmacol 26, 97–106, DOI: 10.1093/ijnp/pyac075.

(62) Preller, K. H., Herdener, M., Pokorny, T., Planzer, A., Kraehenmann, R., Stämpfli, P., Liechti, M. E., Seifritz, E., and Vollenweider, F. X. (2017) The Fabric of Meaning and Subjective Effects in LSD-Induced States Depend on Serotonin 2A Receptor Activation, Curr Biol 27, 451–457, DOI: 10.1016/j.cub.2016.12.030.

(63) Vollenweider, F. X., Vollenweider-Scherpenhuyzen, M. F., Bäbler, A., Vogel, H., and Hell, D. (1998) Psilocybin induces schizophrenia-like psychosis in humans via a serotonin-2 agonist action, Neuroreport 9, 3897–3902, DOI: 10.1097/00001756-199812010-00024.

(64) Brandt, S. D., Kavanagh, P. V., Twamley, B., Westphal, F., Elliott, S. P., Wallach, J., Stratford, A., Klein, L. M., McCorvy, J. D., Nichols, D. E., and Halberstadt, A. L. (2018) Return of the lysergamides. Part IV: Analytical and pharmacological characterization of lysergic acid morpholide (LSM-775), Drug Test Anal 10, 310–322, DOI: 10.1002/dta.2222.

(65) Warren, A. L., Lankri, D., Cunningham, M. J., Serrano, I. C., Parise, L. F., Kruegel, A. C., Duggan, P., Zilberg, G., Capper, M. J., Havel, V., Russo, S. J., Sames, D., and Wacker, D. (2024) Structural pharmacology and therapeutic potential of 5-methoxytryptamines, Nature, DOI: 10.1038/s41586-024-07403-2.

(66) Pokorny, T., Preller, K. H., Kraehenmann, R., and Vollenweider, F. X. (2016) Modulatory effect of the 5-HT1A agonist buspirone and the mixed non-hallucinogenic 5-HT1A/2A agonist ergotamine on psilocybin-induced psychedelic experience, Eur Neuropsychopharmacol 26, 756–766, DOI: 10.1016/j.euroneuro.2016.01.005.

(67) Gattuso, J. J., Wilson, C., Li, S., Hannan, A. J., and Renoir, T. (2025) Mice lacking the serotonin transporter do not respond to the behavioural effects of psilocybin, European Journal of Pharmacology 991, 177304, DOI: 10.1016/j.ejphar.2025.177304.

(68) Becker, A. M., Humbert-Droz, M., Mueller, L., Jelusic, A., Tolev, A., Straumann, I., Avedisian, I., Erne, L., Thomann, J., Luethi, D., Grunblatt, E., Meyer Zu Schwabedissen, H., and Liechti, M. E. (2025) Acute Effects and Pharmacokinetics of LSD after Paroxetine or Placebo Pre-Administration in a Randomized, Double-Blind, Cross-Over Phase I Trial, Clin Pharmacol Ther n/a, DOI: 10.1002/cpt.3618.

(69) Krall, C. M., Richards, J. B., Rabin, R. A., and Winter, J. C. (2008) Marked decrease of LSD-induced stimulus control in serotonin transporter knockout mice, Pharmacology Biochemistry and Behavior 88, 349–357, DOI: 10.1016/j.pbb.2007.09.006.

(70) Qu, Y., Villacreses, N., Murphy, D. L., and Rapoport, S. I. (2005) 5-HT2A/2C receptor signaling via phospholipase A2 and arachidonic acid is attenuated in mice lacking the serotonin reuptake transporter, Psychopharmacology 180, 12–20, DOI: 10.1007/s00213-005-2231-5.

(71) Glatfelter, G., Walther, D., Partilla, J., Chadeayne, A. R., Manke, D. R., and Baumann, M. H. (2024) Pharmacological profiles and psychedelic-like effects of 4-hydroxy-, 4-acetoxy-, and 4-methoxy-N-methyl-N-isopropyltryptamine, The Journal of Pharmacology and Experimental Therapeutics 389, 281, DOI: 10.1124/jpet.281.923160.

(72) Gessner, P. K., Khairallah, P. A., McIsaac, W. M., and Page, I. H. (1960) THE RELATIONSHIP BETWEEN THE METABOLIC FATE AND PHARMACOLOGICAL ACTIONS OF SEROTONIN, BUFOTENINE AND PSILOCYBIN, The Journal of Pharmacology and Experimental Therapeutics 130, 126–133, DOI: 10.1016/S0022-3565(25)25873-6.

(73) Toshihiro, T., Kazuhiro, T., Tatsuo, I., Kazuhiko, Y., Ren, I., Kiichi, I., and Shigeo, N. (1985) 11C-labelling of indolealkylamine alkaloids and the comparative study of their tissue distributions, The International Journal of Applied Radiation and Isotopes 36, 965–969, DOI: 10.1016/0020-708X(85)90257-1.

(74) Silva, M. T. A., and Calil, H. M. (1975) Screening hallucinogenic drugs: Systematic study of three behavioral tests, Psychopharmacologia 42, 163–171, DOI: 10.1007/BF00429548.

(75) McGriff, S. A., Hecker, J. C., Maitland, A. D., Partilla, J. S., Baumann, M. H., and Glatfelter, G. C. (2025) Psychedelic-like effects induced by 2,5-dimethoxy-4-iodoamphetamine, lysergic acid diethylamide, and psilocybin in male and female C57BL/6J mice, Psychopharmacology, DOI: 10.1007/s00213-025-06795-x.

(76) Klein, A. K., Chatha, M., Laskowski, L. J., Anderson, E. I., Brandt, S. D., Chapman, S. J., McCorvy, J. D., and Halberstadt, A. L. (2021) Investigation of the Structure-Activity Relationships of Psilocybin Analogues, ACS Pharmacol Transl Sci 4, 533–542, DOI: 10.1021/acsptsci.0c00176.

(77) Tylš, F., Pálenícek, T., Kaderábek, L., Lipski, M., Kubešová, A., and Horácek, J. (2016) Sex differences and serotonergic mechanisms in the behavioural effects of psilocin, Behavioural Pharmacology 27, 309–320, DOI: 10.1097/fbp.0000000000000198.

(78) Pego, A. M. F., Schoffner, M., Sammeta, V. R., Naeem, M., Manke, D. R., Chadeayne, A., Glatfelter, G. C., Baumann, M. H., and Concheiro-Guisán, M. (2025) Development and validation of an analytical method for the determination of select 4-position ring-substituted tryptamines in plasma by liquid chromatography-tandem mass spectrometry, Journal of Analytical Toxicology, bkaf045, DOI: 10.1093/jat/bkaf045.

(79) Sherwood, A. M., Halberstadt, A. L., Klein, A. K., McCorvy, J. D., Kaylo, K. W., Kargbo, R. B., and Meisenheimer, P. (2020) Synthesis and Biological Evaluation of Tryptamines Found in Hallucinogenic Mushrooms: Norbaeocystin, Baeocystin, Norpsilocin, and Aeruginascin, J Nat Prod 83, 461–467, DOI: 10.1021/acs.jnatprod.9b01061.

(80) Kane, G. A., Lopes, G., Saunders, J. L., Mathis, A., and Mathis, M. W. (2020) Real-time, low-latency closed-loop feedback using markerless posture tracking, eLife 9, e61909, DOI: 10.7554/eLife.61909.

(81) Lapp, H. E., Salazar, M. G., and Champagne, F. A. (2023) Automated maternal behavior during early life in rodents (AMBER) pipeline, Scientific Reports 13, 18277, DOI: 10.1038/s41598-023-45495-4.

(82) Popik, P., Cyrano, E., Piotrowska, D., Holuj, M., Golebiowska, J., Malikowska-Racia, N., Potasiewicz, A., and Nikiforuk, A. (2024) Effects of ketamine on rat social behavior as analyzed by DeepLabCut and SimBA deep learning algorithms, Frontiers in Pharmacology 14, DOI: 10.3389/fphar.2023.1329424.

(83) Halberstadt, A. L., Koedood, L., Powell, S. B., and Geyer, M. A. (2011) Differential contributions of serotonin receptors to the behavioral effects of indoleamine hallucinogens in mice, J Psychopharmacol 25, 1548–1561, DOI: 10.1177/0269881110388326.

(84) Halberstadt, A. L., and Geyer, M. A. (2011) Multiple receptors contribute to the behavioral effects of indoleamine hallucinogens, Neuropharmacology 61, 364–381, DOI: 10.1016/j.neuropharm.2011.01.017.

(85) Halberstadt, A. L. (2015) Recent advances in the neuropsychopharmacology of serotonergic hallucinogens, Behav Brain Res 277, 99–120, DOI: 10.1016/j.bbr.2014.07.016.

(86) Tillmann, J. F., Hsu, A. I., Schwarz, M. K., and Yttri, E. A. (2024) A-SOiD, an active-learning platform for expert-guided, data-efficient discovery of behavior, Nature Methods 21, 703–711, DOI: 10.1038/s41592-024-02200-1.

(87) Ye, S., Filippova, A., Lauer, J., Schneider, S., Vidal, M., Qiu, T., Mathis, A., and Mathis, M. W. (2024) SuperAnimal pretrained pose estimation models for behavioral analysis, Nature Communications 15, 5165, DOI: 10.1038/s41467-024-48792-2.

(88) Kapoor, S., and Narayanan, A. (2023) Leakage and the reproducibility crisis in machine-learning-based science, Patterns 4, 100804, DOI: 10.1016/j.patter.2023.100804.

(89) Luethi, D., Glatfelter, G. C., Pottie, E., Sellitti, F., Maitland, A. D., Gonzalez, N. R., Kryszak, L., Jackson, S., Hoener, M. C., Stove, C. P., Liechti, M. E., Smiesko, M., Baumann, M. H., Simmler, L. D., and Rudin, D. (**In review**) The 4-alkyl chain length of 2,5-dimethoxyamphetamines differentially affects in vitro serotonin receptor actions versus in vivo psychedelic-like effects, Mol Psychiatry.

(90) Insafutdinov, E., Pishchulin, L., Andres, B., Andriluka, M., and Schiele, B. (2016) DeeperCut: A Deeper, Stronger, and Faster Multi-person Pose Estimation Model, In Computer Vision – ECCV 2016 (Leibe, B., Matas, J., Sebe, N., and Welling, M., Eds.), pp 34–50, Springer International Publishing, Cham.

(91) He, K., Zhang, X., Ren, S., and Sun, J. (2015) Deep Residual Learning for Image Recognition, *arXiv [cs.CV]*.

(92) Glatfelter, G. C., Partilla, J. S., and Baumann, M. H. (2022) Structure-activity relationships for 5F-MDMB-PICA and its 5F-pentylindole analogs to induce cannabinoid-like effects in mice, Neuropsychopharmacology 47, 924–932, DOI: 10.1038/s41386-021-01227-8.

(93) Schindler, C. W., Thorndike, E. B., Partilla, J. S., Rice, K. C., and Baumann, M. H. (2021) Amphetamine-like Neurochemical and Cardiovascular Effects of α-Ethylphenethylamine Analogs Found in Dietary Supplements, J Pharmacol Exp Ther 376, 118–126, DOI: 10.1124/jpet.120.000129.

(94) Glatfelter, G. C., Walther, D., Evans-Brown, M., and Baumann, M. H. (2021) Eutylone and Its Structural Isomers Interact with Monoamine Transporters and Induce Locomotor Stimulation, ACS Chemical Neuroscience 12, 1170–1177, DOI: 10.1021/acschemneuro.0c00797.

(95) Solis, E., Jr., Partilla, J. S., Sakloth, F., Ruchala, I., Schwienteck, K. L., De Felice, L. J., Eltit, J. M., Glennon, R. A., Negus, S. S., and Baumann, M. H. (2017) N-Alkylated Analogs of 4-Methylamphetamine (4-MA) Differentially Affect Monoamine Transporters and Abuse Liability, Neuropsychopharmacology 42, 1950–1961, DOI: 10.1038/npp.2017.98.

