## Supporting Information File - Tables & Figures for "Rapid, open-source, and automated quantification of the head twitch response in C57BL/6J mice using DeepLabCut and Simple Behavioral Analysis"

**Table of contents:**

| Figure/Table | Page # |
|---|---|
| Final snapshot table for the DLC model at each video resolution (**Table S1**); | **S2** |
| DLC inference time and FFmpeg downscaling time cost table (**Table S2**); | **S2** |
| DLC pose estimation model time costs for 50% resolution (906 x 680); 120 fps videos (**Table S3**); | **S3** |
| SimBA behavioral classifier model training time in minutes for 50% resolution (906 x 680), 120 fps model (**Table S4**); | **S3** |
| FFmpeg video downscaling time, DLC inference time, and SimBA video analysis for HTR quantification times in minutes for 50% resolution; 120 fps model combination at 0.25 discrimination threshold (**Table S5**); | **S4** |
| heatmap plots of accuracy-discrimination threshold curve summary data (**Fig. S1**); | **S5** |
| accuracy-discrimination threshold curves from a second experimenter (**Fig. S2**); | **S6** |
| heatmap plots of accuracy-discrimination threshold curve summary data from a second experimenter (**Fig. S3**); | **S7** |
| time-course plots for acute effects of bufotenine on the HTR (**Fig. S4**); | **S8** |
| time-course plots for acute effects of bufotenine on motor activity (**Fig. S5**); | **S8** |
| summary statistics table for acute effects of bufotenine (**Table S6**); | **S9** |
| summary statistics table for further validation studies (**Table S7**); | **S10** |
| heatmap plots illustrating monoamine transporter activity of bufotenine (**Fig. S6**) | **S10** |

**Table S1**. Final snapshot 500,000 for shuffle 3 for each DLC model (computed root mean square error between the model-predicted labels and user labels). Comparisons standardized using % of mouse body length (measured from nose to tail base in representative frame from experimental videos).

| Video resolution | Train error (pixels) | Test error (pixels) | % of mouse body length |
|---|---|---|---|
| **50%** | 1.76 | 2.92 | 1.82 |
| **25%** | 2.22 | 2.87 | 2.54 |
| **12.5%** | 1.27 | 1.87 | 2.33 |

**Table S2**. FFmpeg video downscaling time, and DLC inference time in minutes for a subset of 10 training videos across different video resolutions on a A100 Nvidia GPU. Ten videos were 168 minutes combined with 13 body parts identified.

| | **120 fps** | | **60 fps** | | **30 fps** | |
|---|---|---|---|---|---|---|
| | **FFmpeg video downscaling time** | **DLC inference time** | **FFmpeg video downscaling time** | **DLC inference time** | **FFmpeg video downscaling time** | **DLC inference time** |
| **50%** | 92.70 | 342.45 | 47.35 | 189.12 | 32.47 | 91.60 |
| **25%** | 58.68 | 179.73 | 34.43 | 91.37 | 27.77 | 45.45 |
| **12%** | 45.58 | 97.95 | 29.45 | 57.75 | 26.12 | 28.57 |

**Table S3**. DLC pose estimation model time costs in minutes for 50% resolution (906 x 680), 120 fps videos. Time costs listed are for one shuffle featuring 10 historical experimental videos (144 min). Pose estimation model was trained for three shuffles (500,000 iterations per shuffle) with batch size = 8. In total, 30 historical experimental videos (415 min) were used for training DLC pose estimation model.

| | |
|---|---|
| Create new project | Less than 1s |
| Extract outlier frames (20 frames per video, k-means clustering) | 78.53 |
| Manual labeling of 13 body parts for 200 frames | 68.88 |
| Check labels | 1.08 |
| Create training set | Less than 1s |
| Train model (500k iterations) | 583.58 |
| Analyze new videos (10 videos) | 323.85 |
| **Total** | 1,055.92 |

**Table S4**. SimBA behavioral classifier model training time in minutes for 50% resolution (906 x 680), 120 fps model. Ten historical experimental videos (168 min) were used for training SimBA behavioral classifier. Videos were imported as symbolic links.

| | |
|---|---|
| Import 10 videos | Less than 1s |
| Import 10 .csv files | 2.35 |
| Standardize distance (pixels/mm) | 0.98 |
| Skip outlier correction | 1.05 |
| Extract features | 0.89 |
| Manually label 136 HTR events | 65.95 |
| Train behavioral classifier | 34.99 |
| **Total** | 105.91 |

**Table S5.** FFmpeg video downscaling time, DLC inference time, and SimBA video analysis for HTR quantification times in minutes for 50% resolution, 120 fps model combination at 0.25 discrimination threshold. Forty-seven new experimental videos (692 min) were analyzed for a single bufotenine experiment using 50%, 120 fps model combination. Videos were imported as symbolic links.

| | | |
|---|---|---|
| FFmpeg | Video downscaling to 50% (906 x 680), 120 fps | 380.90 |
| DLC | Analyze 47 videos using 50% model | 1405.70 |
| SimBA | Import 47 videos | Less than 1s |
| | Import 47 .csv files | 9.53 |
| | Standardize distance (pixels/mm) | 4.37 |
| | Skip outlier correction | 4.24 |
| | Extract features | 3.67 |
| | Run machine model | 6.25 |
| | Analyze machine predictions (aggregates) | 0.23 |
| **Total** | | 1,814.89 |

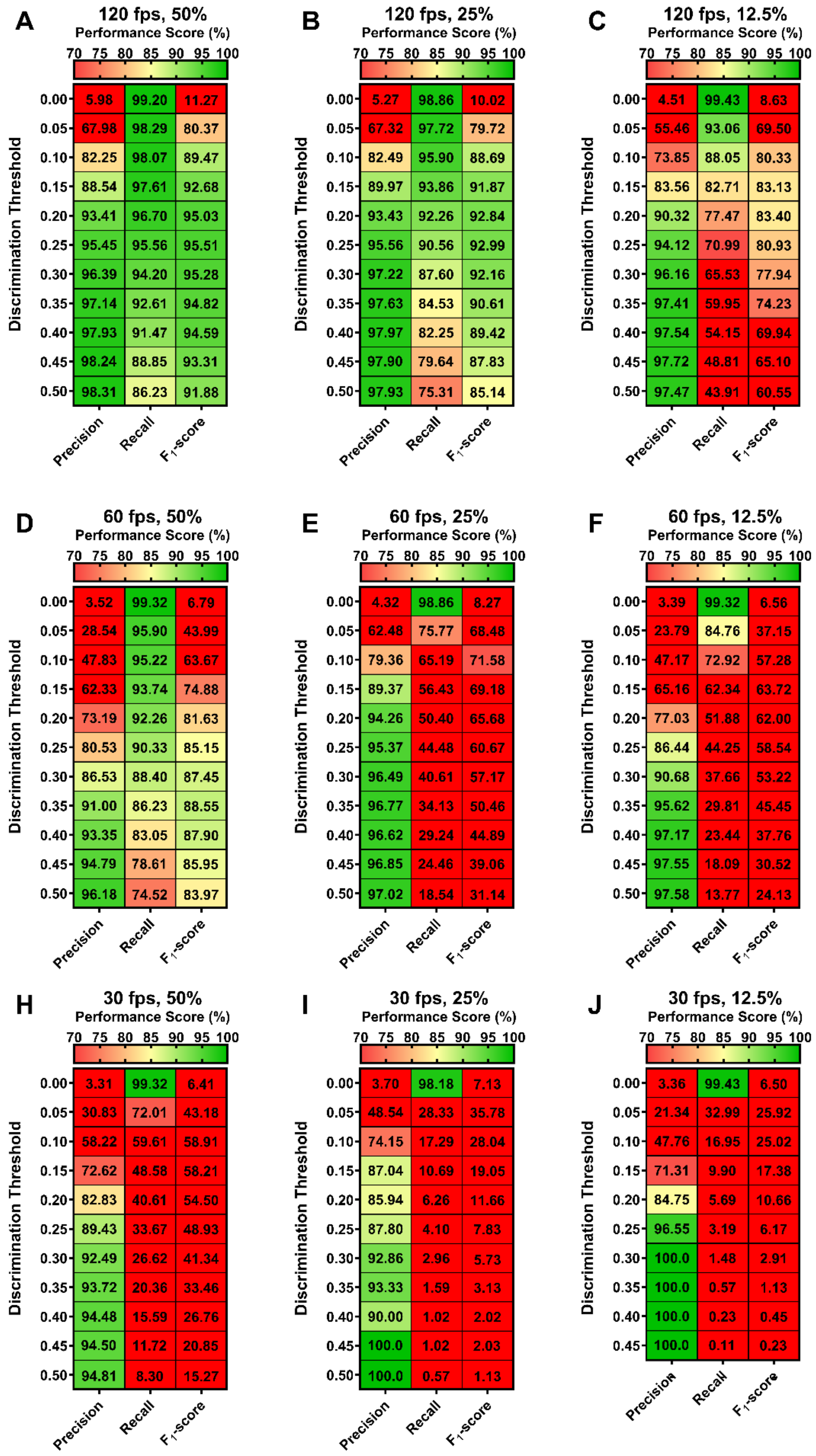


**Figure S1**. Heatmap visualization summarizing accuracy-discrimination threshold curves for SimBA behavioral classifiers for 50% (**A**, **D**, **H**), 25% (**B**, **E**, **I**), and 12.5% (**D**, **F**, **J**) video resolution at 120 (**A** – **C**), 60 (**D** – **F**), and 30 fps ( **H** – **J**). All minimum behavior bout lengths were set to 30 ms.

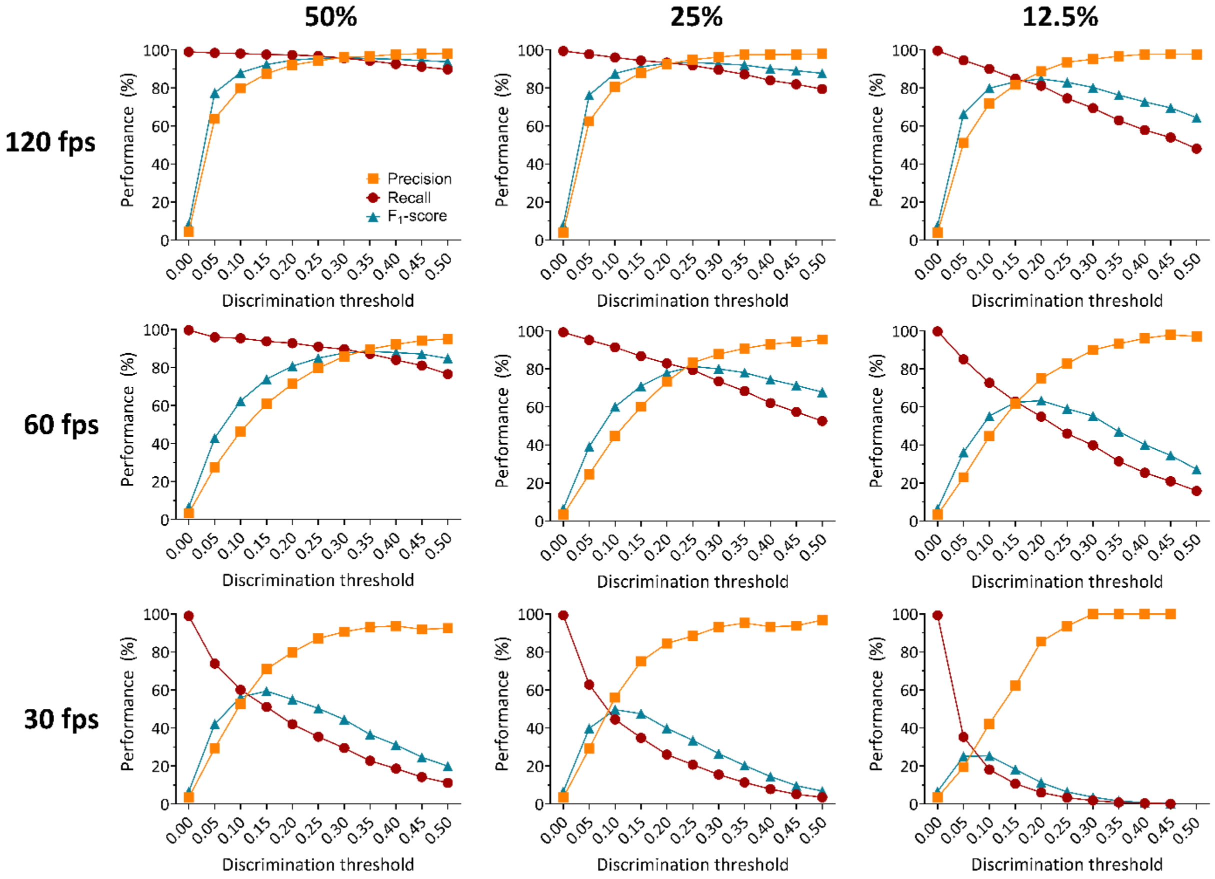


**Figure S2.** Accuracy-discrimination threshold curves for SimBA classifiers for 50%, 25%, and 12.5% video resolution at 120, 60, and 30 fps as determined by another experimenter in the lab. Discrimination thresholds are the probability levels at which the behavioral classifier determines an HTR is detected. All minimum behavior bout lengths were set to 30 ms. Precision in red, recall in orange, and $F_1$ scores in teal at different discrimination thresholds are shown for all behavioral classifiers.

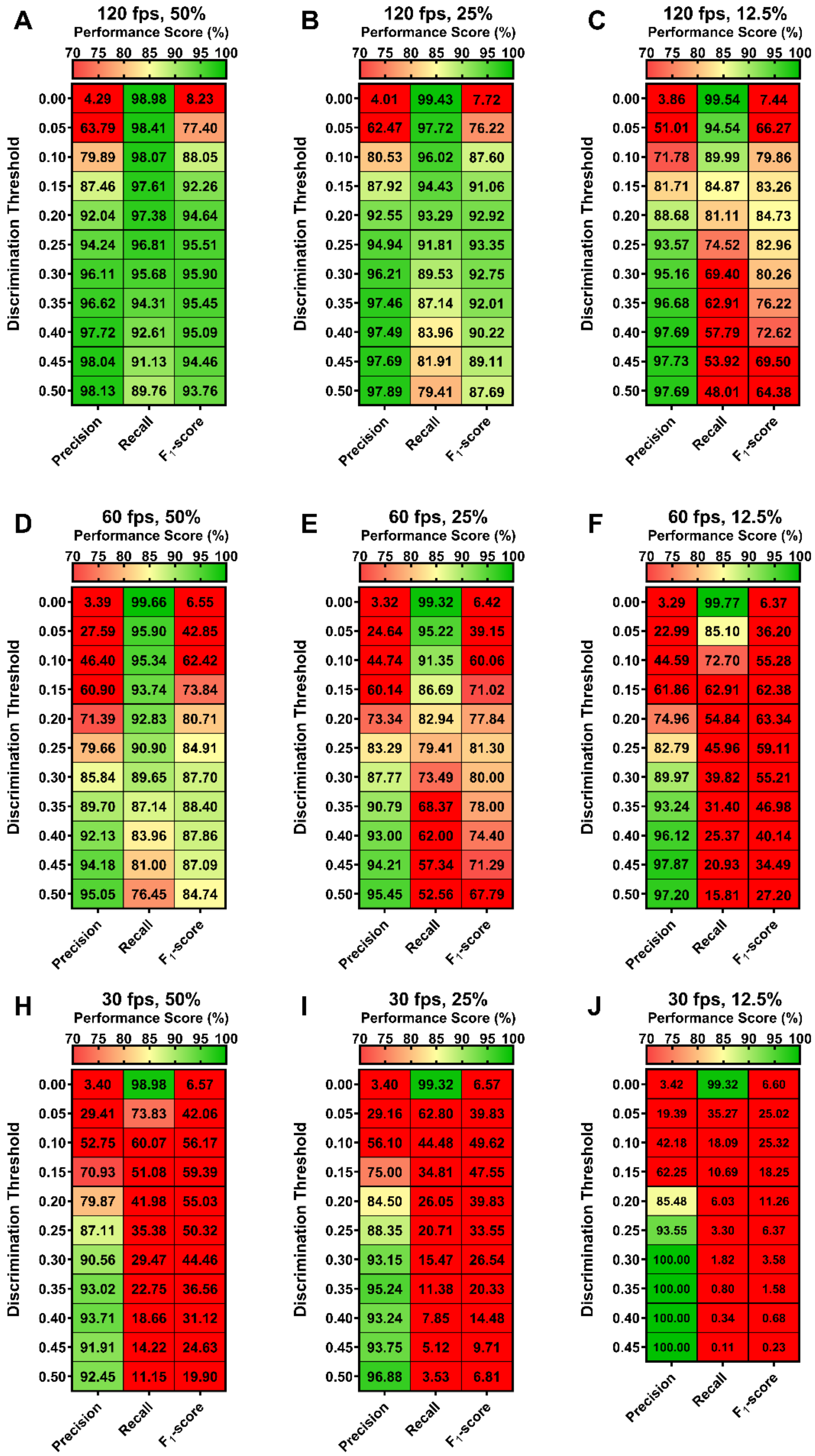


**Figure S3**. Heatmap visualization summarizing accuracy-discrimination threshold curves for SimBA behavioral classifiers as determined by a second experimenter for 50% (**A**, **D**, **H**), 25% (**B**, **E**, **I**), and 12.5% (**D**, **F**, **J**) video resolution at 120 (**A** – **C**), 60 (**D** – **F**), and 30 fps ( **H** – **J**). All minimum behavior bout lengths were set to 30 ms.

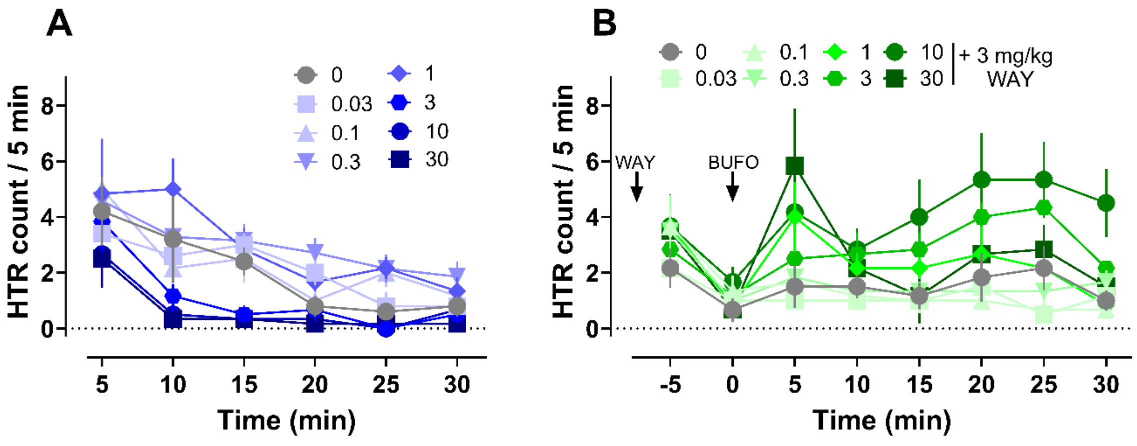


**Figure S4**. Time-course plots for effects of bufotenine without (**A**) and with WAY100635 pretreatment (**B**) on HTR during the 30 min testing period. All values are mean ± SEM and represent *n* = 5 – 7 mice per data point.

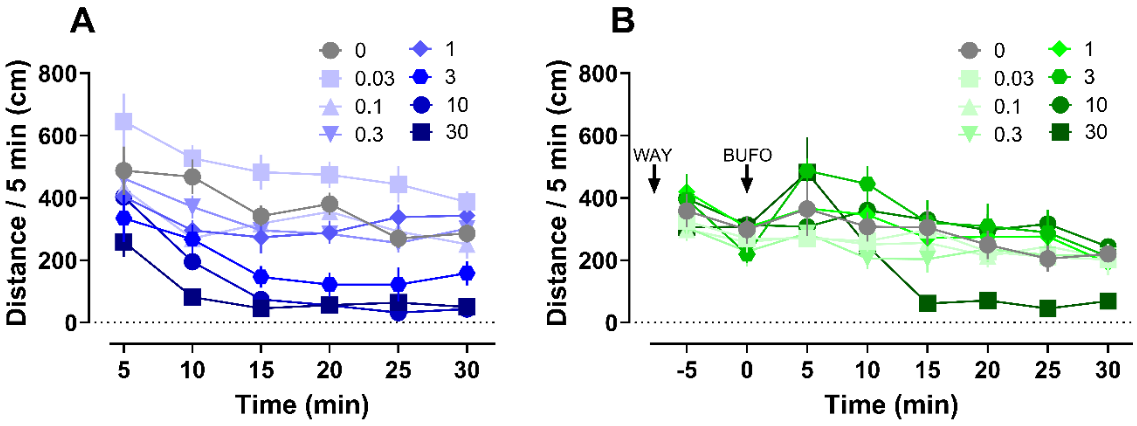


**Figure S5**. Time-course plots for effects of bufotenine without (**A**) and with WAY100635 pretreatment (**B**) on motor activity during the 30 min testing period. All values are mean ± SEM and represent *n* = 5 – 7 mice per data point.

**Table S6.** Mean ± SEM and post hoc test comparisons vs. vehicle controls for dose-response effects of 5-HO-DMT without and with WAY100635 (3 mg/kg) pretreatment. Bold values are statistically significant values vs. saline vehicle control (0 mg/kg, Dunnett's or Dunnett's T3) or differences between doses (dose-response vs. WAY experiments, Sidak's). Additional statistical information can be found in the **Materials and Methods** section. The overall ANOVA values for experiments without and with WAY pretreatment were: HTR (F (7, 38) = 5.643 $p$ = 0.0002 & F (7, 40) = 5.306 $p$ = 0.0002), temperature change (W = 52.26 (7.000, 16.66) $p$ < 0.0001 & W = 7.775 (7.000, 16.47) $p$ < 0.0001), and motor activity (F (7, 40) = 13.36 $p$ < 0.0001 & F (7, 40) = 2.973 $p$ = 0.0132). Overall two-way ANOVA $F$-test values for HTR, temperature change, and locomotor activity data set comparisons respectively were: HTR-dose (F (7, 78) = 1.716 $p$ = 0.1174, HTR-treatment F (1, 78) = 2.205 $p$ = 0.1416, HTR-interaction F (7, 78) = 8.913 $p$ < 0.0001, temp-dose (F (7, 80) = 115.8 $p$ < 0.0001, temp-treatment F (1, 80) = 54.00 $p$ < 0.0001, temp-interaction F (7, 80) = 19.78 $p$ < 0.0001, motor-dose (F (7, 80) = 8.918 $p$ < 0.0001, motor-treatment F (1, 80) = 1.477 $p$ = 0.2278, and motor-interaction F (7, 80) = 8.850 $p$ < 0.0001.

| **Drug *n* per dose** | **Dose** | **HTR (total events)** | | **vs. 0 mg/kg Post Test HTR** | **Body Temp Δ (°C)** | | **vs. 0 mg/kg Post Test Temp** | **Locomotor Activity (cm)** | | **vs. 0 mg/kg Post Test Motor** | **Dose by dose vs. WAY Post Test HTR** | **Post Test Temp** | **Post Test Motor** |
|---|---|---|---|---|---|---|---|---|---|---|---|---|---|
| | **(mg/kg)** | **Mean** | **SEM** | ***p* value** | **Mean** | **SEM** | ***p* value** | **Mean** | **SEM** | ***p* value** | ***p* value** | ***p* value** | ***p* value** |
| 5-HO-DMT *n* = 5 - 7 | 0 | 12 | 3.1 | - | -0.16 | 0.07 | - | 2233 | 214 | - | >0.9999 | >0.9999 | 0.7232 |
| | 0.03 | 12.7 | 1.4 | >0.9999 | 0.00 | 0.07 | 0.6176 | 2959 | 292 | 0.4042 | 0.9329 | >0.9999 | <0.0001 |
| | 0.1 | 13.8 | 2.6 | >0.9999 | -0.07 | 0.11 | 0.9846 | 1915 | 192 | 0.9995 | 0.6781 | >0.9999 | 0.9170 |
| | 0.3 | 17.7 | 2.6 | 0.9704 | -0.26 | 0.12 | 0.9843 | 1971 | 227 | >0.9999 | 0.3474 | >0.9999 | 0.4911 |
| | 1 | 17.8 | 2.6 | 0.9741 | -0.12 | 0.05 | 0.9978 | 1942 | 173 | 0.9999 | 0.9998 | >0.9999 | >0.9999 |
| | 3 | 6.6 | 2.5 | 0.9935 | **-2.11** | **0.26** | **0.0010** | **1150** | **258** | **0.0080** | 0.0899 | **<0.0001** | **0.0313** |
| | 10 | 4.5 | 2.2 | 0.7843 | **-4.98** | **0.52** | **0.0014** | **803** | **147** | **0.0002** | **<0.0001** | **<0.0001** | **0.0098** |
| | 30 | 3.7 | 1.6 | 0.6163 | **-6.37** | **0.36** | **<0.0001** | **557** | **132** | **<0.0001** | **0.0376** | **<0.0001** | 0.9721 |
| WAY100635 (3mg/kg) + 5-HO-DMT *n* = 6 | 0 | 9.2 | 2.6 | - | -0.23 | 0.15 | - | 1651 | 240 | - | - | - | - |
| | 0.03 | 6.2 | 1.7 | >0.9999 | -0.20 | 0.04 | >0.9999 | 1468 | 130 | >0.9999 | - | - | - |
| | 0.1 | 6.1 | 1.5 | >0.9999 | -0.25 | 0.09 | >0.9999 | 1448 | 186 | >0.9999 | - | - | - |
| | 0.3 | 8.8 | 0.8 | >0.9999 | -0.37 | 0.13 | 0.9898 | 1362 | 169 | 0.9998 | - | - | - |
| | 1 | 14 | 2.6 | 0.9949 | -0.18 | 0.07 | >0.9999 | 1724 | 77.2 | >0.9999 | - | - | - |
| | 3 | 18.5 | 2.9 | 0.3262 | -0.45 | 0.11 | 0.8597 | 2062 | 308 | 0.9760 | - | - | - |
| | 10 | **26.2** | **5.4** | **0.0007** | -1.12 | 0.22 | 0.0540 | 1853 | 180 | >0.9999 | - | - | - |
| | 30 | 16.2 | 4.0 | 0.8123 | **-4.03** | **0.56** | **0.0036** | 973.3 | 167 | 0.3651 | - | - | - |

**Table S7.** Mean ± SEM and post hoc test comparisons vs. vehicle controls (Dunnett's T3) for HTRs induced by saline vehicle, psilocybin, DOI, or LSD. Bold values ae statistically significant values vs. saline vehicle control (0 mg/kg). Additional statistical information can be found in the **Materials and Methods** section. The overall Welch's ANOVA value for HTR was: W = 52.74 (3.000, 9.391) $p < 0.0001$.

.

| Treatment | *n* per group | Dose | HTR (total events) | | vs. 0 mg/kg Post Test HTR |
|---|---|---|---|---|---|
| | | (mg/kg) | Mean | SEM | *p* value |
| Vehicle | 6 | 0 | 8.8 | 2.2 | - |
| Psilocybin | 6 | 1 | **34.0** | **4.2** | **0.0030** |
| DOI | 5 | 1 | **119.8** | **14.4** | **0.0041** |
| LSD | 6 | 0.1 | **52.3** | **3.1** | **<0.0001** |

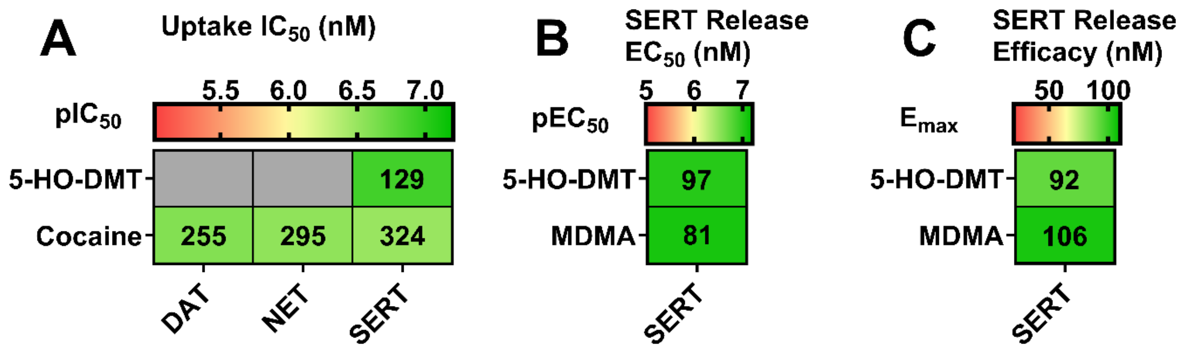


**Figure S6.** Potency of bufotenine (5-HO-DMT) for uptake (**A**) and release (**B**, **C**) at monoamine transporters (dopamine transporter or DAT, norepinephrine transporter or NET, and SERT) in rat brain synaptosomes. Data are from $n$ = 3 experiments performed in triplicate, as determined from nonlinear regression of the concentration-response curves.
